# *CarbMetSim:* A discrete-event simulator for carbohydrate metabolism in humans

**DOI:** 10.1101/491019

**Authors:** Mukul Goyal, Buket Aydas, Husam Ghazaleh, Sanjay Rajasekharan

## Abstract

This paper describes *CarbMetSim*, a *discrete-event* simulator that tracks the blood glucose level of a person in response to a timed sequence of diet and exercise activities. *CarbMetSim* implements broader aspects of carbohydrate metabolism in human beings with the objective of capturing the average impact of various diet/exercise activities on the blood glucose level. Key organs (stomach, intestine, portal vein, liver, kidney, muscles, adipose tissue, brain and heart) are implemented to the extent necessary to capture their impact on the production and consumption of glucose. Key metabolic pathways (*glucose oxidation, glycolysis* and *gluconeogenesis*) are accounted for in the operation of different organs. The impact of insulin and insulin resistance on the operation of various organs and pathways is captured in accordance with published research. *CarbMetSim* provides broad flexibility to configure the insulin production ability, the average flux along various metabolic pathways and the impact of insulin resistance on different aspects of carbohydrate metabolism. The simulator does not yet have a detailed implementation of protein and lipid metabolism.

## 1 Introduction

More than 400 million people world wide suffer from diabetes [1]. People with *Type 2 Diabetes* (around 90% of total diabetic population [1]) usually have at least some ability to produce insulin, however their bodies develop *insulin resistance* and hence are not able to react strongly enough to the presence of insulin in blood to keep the *blood glucose level* (BGL) under control. On the other hand, people with *Type 1 Diabetes* cannot produce insulin endogenously at all and hence must receive external insulin regularly. Keeping BGL under control is a constant struggle for people with diabetes. One wrong meal choice may result in very high BGL and an accompanying feeling of sickness for several hours. Persistently high BGL would ultimately cause a number of severe complications such as heart/kidney failure, blindness and limb amputations. Those using external insulin may suffer life threatening hypoglycemic incidents if too much insulin is injected. Physical exercise allows the muscles to use glucose in the blood even in the absence of insulin but exercise activities need to be carefully coordinated with food and medication intake so as to avoid hypoglycemia. For people with Type 1 Diabetes, physical exercise may even worsen the state of hyperglycemia. In general, people with diabetes need help deciding how they should plan their food and exercise activities so as to keep their BGL under control. There is a real need for tools that help diabetic people understand the impact a particular sequence of food and exercise activities would have on their BGL. Continuous BGL monitoring solutions, now offered by a number of vendors, can significantly help but are either not easily available to a vast majority of diabetic people world-wide or are simply too expensive. Clearly, one solution is to build simulation tools that use our vast knowledge of energy metabolism in human beings to give reasonably accurate prediction of the impact of a diet/exercise sequence on some one’s BGL. A few such simulators already exist [2, 3] but are geared towards predicting the impact of *individual* meals and are not available in a format that can be freely used by individuals. This paper describes *CarbMetSim* (the **Carb**ohydrate **Met**abolism **Sim**ulator), an *open-source* and freely available [4] simulation software that predicts minute by minute BGL in response to an arbitrary length sequence of food and exercise activities. While the existing simulation tools are based on *continuous time* models that use differential and algebraic equations to describe physiological details, *CarbMetSim* is based on a *discrete event* model where the time increments in units (called *ticks*) one minute long. At the beginning of each tick, *CarbMetSim* fires the food/exercise events that need to be fired at this time and directs various simulated body organs to do the work they are supposed to do during this tick. The simulator is currently geared for use by people with Prediabetes and Type 2 Diabetes who follow a fixed medication (including long term insulin) regime prescribed by their physicians. Future versions of the simulator will include the ability to specify the dosage of externally injected short term insulin to allow use by people dependent on short term insulin (including those with Type 1 Diabetes).

*CarbMetSim* implements broader aspects of carbohydrate metabolism in human beings with the objective of capturing the average impact of various diet/exercise activities on the BGL of people with different levels of diabetes. The simulator implements key organs (stomach, intestine, portal vein, liver, kidney, muscles, adipose tissue, brain and heart) to the extent necessary to capture their impact on the production and consumption of glucose. Key metabolic pathways (*glucose oxidation, glycolysis* and *gluconeogenesis*) are accounted for in the operation of different organs. The impact of insulin and insulin resistance on the operation of various organs/pathways is captured in accordance with published research. *CarbMetSim* provides broad flexibility to configure the insulin production ability, the average flux along various metabolic pathways and the impact of insulin resistance on different aspects of carbohydrate metabolism. Thus, it is possible to customize the simulator for a particular user by setting appropriate values to various configurable parameters.

*CarbMetSim* is not yet a finished product. The protein and lipid metabolism are implemented in a very simplified manner. The simulator does not yet consider monosaccharides other than glucose and assumes that all dietary carbohydrate gets converted to glucose after digestion. The impact of insulin is captured in a simplified manner and other important hormones (e.g. glucagon) are not yet directly modeled. Impact of externally injected short term insulin is not modeled yet. Only aerobic exercise activities can be simulated at present. Finally, *CarbMetSim* is not yet capable of translating a user’s diet/exercise/BGL data into the values of simulation parameters governing the behavior of different organs. The simulator has broad applicability beyond its original purpose described above. For example, it is possible to extend the implementation to study the long term impact of diabetes on various organs or to predict changes in body weight in response to a diet/exercise regimen.

## 2 Modeling Carbohydrate Metabolism in Humans - A Literature Review

Existing approaches to model carbohydrate metabolism in human beings can be classified as either *data-driven* or *knowledge-driven* [5]. *The data-driven (or empirical*) models relate a user’s recent BGL values along with other relevant information (e.g. diet, exercise, medication and stress) to the user’s future BGL values using approaches such as neural networks [6–11] and gaussian models [12]. A number of different neural network models exist including those based on multilayer perceptrons [7, 8], radial basis function [9], wavelets [10], time series convolution [6] and recurrent neural networks [6, 11]. Such models consider the human body to be a *black-box* and do not take in account the physiological aspects of carbohydrate metabolism [13].

Unlike the data-driven models, the knowledge-driven models are based on human physiology. In such models, different factors are treated as different *compartments* that influence each other and are described by a set of differential and algebraic equations [5]. The earliest such models [14–16] involved two *linear* compartments - one for glucose in blood and the other for insulin in blood - such that the rates of appearance/disappearance of glucose/insulin were *linearly* proportional to their level in blood. The next generation of models included *non-linear* rates and consideration of additional hormones (e.g. glucagon) besides insulin [17]. Foster [18] presented a six compartment model, one each for blood glucose, liver glycogen, muscle glycogen, plasma insulin, plasma glucagon and free fatty acids in plasma, and the addition/removal from each compartment happened in a non-linear fashion. Some of the other notable multi-compartment, nonlinear models were those developed by Cerasi [19], Insel [20], Cramp and Carson [21] and Cobelli et al. [22]. These models were increasingly more complex with many physiological details taken in account. Sorensen [23] provides a good overview of the earliest knowledge-based models (and some of the later models described next).

Bergman et al. [24, 25] designed a method to quantify a) the sensitiviy of an individual’s beta cells to his/her BGL and b) the sensitivity of the individual’s BGL to insulin level in his/her blood. For this purpose, a *minimally* complex mathematical model was developed that could capture the individual differences in two sensitivities mentioned above. Bergman’s *minimal* model has been modified in a variety of ways. Furler et al. [26] introduced modifications to allow for absence of insulin production by pancreas and external insulin infusion. Bergman’s model has also been used to study closed [27] and semi-closed [28] loop optimal control algorithms to determine the insulin infusion profile for an individual. Roy and Parker extended Bergman’s model to take in account the level of *free fatty acids* in plasma [29]. Bergman’s model has also been extended to take in account the impact of physical exercise [30, 31].

Tiran et al. [32] developed a multi-compartment model for glucose circulation where each relevant organ was modeled as a separate compartment. Guyton et al. [33] developed another multi-compartment model consisting of a glucose circulation subsystem (separate compartments for liver glucose, liver glycogen, kidney glucose, brain tissue glucose, brain blood glucose, peripheral (muscles, adipose tissue) blood glucose, peripheral tissue glucose, central (i.e. gastrointestinal tract) blood glucose and central tissue glucose) and an insulin circulation subsystem (separate compartments for liver insulin which represents insulin from pancreatic beta cells, kidney insulin, peripheral blood insulin, peripheral tissue insulin, central blood insulin and central tissue insulin). The model consisted of a total of 32 nonlinear ordinary differential equations (ODEs) with 11 nonlinear ODEs just to model insulin secretion from pancreas [23]. Sorensen [23] presented another physiologically complex, multi-compartment model albeit with a much simplified model for pancreatic insulin secretion. Sorensen’s model consisted of a total of 22 nonlinear ODEs of which 11 ODEs were associated with glucose circulation, 10 ODEs with insulin and 1 ODE with glucagon. Parker et al. [34, 35] updated the Guyton/Sorensen models by accounting for uncertainty in parameter values and by including a model for gastric emptying of carbohydrates in a meal [36]. Hovorka et al. [37] developed a multi-compartment model of glucose and insulin kinetics as part of a model predictive controller for subcutaneous insulin infusion for people with Type 1 Diabetes. This model consists of a two-compartment glucose subsystem (accounting for glucose absorption, distribution and disposal), a two-compartment insulin subsystem (accounting for insulin absorption, distribution and disposal) and an insulin action subsystem (accounting for insulin action on glucose transport, disposal and endogeneous production).

Dalla Man et al. [38] developed a model that related the plasma concentrations of glucose and insulin to various glucose and insulin related rates (the rate of appearance of glucose from the gastro-intestinal tract, the rate at which the glucose is produced by liver and kidney, insulin dependent and independent rates of glucose utilization, the rate of renal extraction of glucose, the rate of insulin secretion by beta cells and the rate of insulin degradation). The parameters of this model were determined using the experimental data collected for 204 normal and 14 Type 2 Diabetic subjects. This model was used to simulate patient behavior in UVA/PADOVA Type 1 Diabetes Simulator [2] aimed at investigating the closed control strategies for insulin pumps. A new version of UVA/PADOVA Type 1 Diabetes Simulator [3] modifies Dalla Man’s model by incorporating glucagon secretion/action/kinetics and nonlinear increase in insulin dependent glucose utlization as BGL dips below the normal range.

The *CarbMetSim* simulator presented in this paper is physiologically complex just like the models presented by Tiran et al. [32], Guyton et al. [33], Sorensen [23] and Dalla Man [38]. The key difference is that *CarbMetSim* implements the physiological details in software with various body organs implemented as *objects* whereas the existing models used ODEs to model physiological details. It can be argued that implementing physiological details in software allows for much more complex behavior to be taken in account than what is possible using ODEs. Moreover, it is much easier to modify physiological behavior implemented in software than via ODEs. In that sense, the presented simulator is an improvement over existing ODE based approaches. It is hoped that these benefits coupled with its *open-source* nature will allow *CarbMetSim* to emerge as a popular simulation model of human metabolism for both diabetes research and self-management tools for diabetic people.

## 3 Key Aspects in *CarbMetSim* Design

In the following, we describe some of the key aspects of *CarbMetSim*’s design.

### 3.1 Food, Exercise and Human Subject Description

In *CarbMetSim*, a food is described in terms of its serving size and the amount of *rapidly available glucose* (RAG), *slowly available glucose* (SAG), protein and fat per serving. The RAG contents include sugars and the *rapidly digestible* starch (i.e. starch that gets digested in vitro within 20 minutes [39, 40]). The SAG contents include the *slowly digestible* starch (i.e. starch that gets digested in vitro between 20 and 120 minutes [39, 40]). In general, the starch with high amylopectin to amylose ratio is classified as rapidly digestible starch whereas the one with high amylose to amylopectin ratio is classified as slowly digestible starch. The *non-starch polysaccharide* (also known as *dietary fiber*) part of the carbohydrates is currently ignored (even though the fiber contents of the food are known to have an impact on the *gastric emptying*).

*CarbMetSim* currently does not have a detailed implementation of the protein and lipid metabolism. However, it does model the impact of protein and fat contents of food on gastric emptying. Hence, the food description should include the total amount of protein and total amount of fat per serving. *CarbMetSim* currently does not characterize protein in terms of its amino acid contents. Since only 3 of the 20 amino acids have *branched chains*, a general assumption is made that 85% of amino acids resulting from protein digestion have *unbranched chains* and the remaining have *branched chains* [41].

*CarbMetSim* can currently simulate only *aerobic* exercises. In *CarbMetSim*, an exercise activity is described in terms of its intensity in units of *Metabolic Equivalent of Task* or *MET*s, where 1 *MET* is 1 *kcal* of energy expenditure per kg of body weight per hour. By convention, 1 *MET* is considered equivalent to 3.5*ml* of oxygen consumption per kg of body weight per minute. Each individual has a certain maximal rate at which he/she can consume oxygen. This individual-specific maximal rate, called *V O*_2_*max*, depends on the gender, age and fitness level of the individual [54]. The intensity of an exercise activity in terms of the associated oxygen consumption rate (described as the %age of the individual’s *V O*_2_*max*, henceforth referred to as %*V O*_2_*max*) determines to a large extent the relative fraction of the glucose and fatty acids oxidized to meet the energy needs of the exercising muscles. Thus, *CarbMetSim* needs to know the gender, age and (self-assessed) fitness level within the age group.of the human subject being simulated. This information is used to estimate the *V O*_2_*max* for the human subject using the tables in Kaminsky et al. [54].

### 3.2 Modeling Insulin Production

The insulin level in blood generally depends on the BGL. If the BGL is high, the insulin level increases as well so as to signal the liver and the muscles to absorb glucose from the blood stream and also to signal both the liver and the kidneys to slow down or stop the endogeneous glucose production via glycogen breakdown and gluconeogenesis. Also, the insulin level in blood decreases in response to physical exercise [65, 73–76] so as to signal the liver and the kidneys to ramp up the endogeneous glucose production. In *CarbMetSim*, the current insulin level in the blood is represented by a variable called *insulinLevel* (inside the *Blood* object) that assumes values between 0 and 1. The value of *insulinLevel* depends on the current BGL, the current exercise intensity (in %*V O*_2_*max*) and a number of configurable parameters: *minGlucoseLevel*_ (typical hypoglycemic BGL), *baseGlucoseLevel*_ (typical fasting BGL), *highGlucoseLevel*_(typical peak BGL) (*minGlucoseLevel*_ *< baseGlucoseLevel*_ *< highGlucoseLevel*_), *baseInsulinLevel*_ (representing the typical fasting insulin level), *peakInsulinLevel*_ (representing typical insulin level when BGL is at peak) (where 0 *≤ baseInsulinLevel*_ *≤ peakInsulinLevel*_ *≤* 1),, *restIntensity*_ (the oxygen consumption rate in %*V O*_2_*max* when the individual is not exercising, by default 2 *MET*s converted to %*V O*_2_*max*) and *intensityPeakGlucoseProd*_ (the exercise intensity in %*V O*_2_*max* at which the liver and kidney produce glucose at the maximum rate, by default 20%). The following rules govern the value of *insulinLevel* :

- If the current BGL is less than or equal to the *minGlucoseLevel*_, the *insulinLevel* stays at value zero.
- If the current BGL is between the *minGlucoseLevel*_ and the *baseGlucoseLevel*_, the *insulinLevel* depends on whether the individual being simulated is currently engaged in exercise or not. If the individual is exercising and
  – if the exercise intensity is greater than or equal to *intensityPeakGlucoseProd*_, the *insulinLevel* stays at zero.
  – otherwise, the *insulinLevel* depends on the exercise intensity. As the exercise intensity decreases from *intensityPeakGlucoseProd* to the the *restIntensity*_, the *insulinLevel* increases linearly from zero to the *baseInsulinLevel*_.

If the individual is not exercising, as the BGL increases from *minGlucoseLevel* to *baseGlucoseLevel*, the *insulinLevel* increases linearly from zero to the *baseInsulinLevel*.

- As the BGL increases from the *baseGlucoseLevel*_ to the *highGlucoseLevel*_, the *insulinLevel*_ increases linearly from the *baseInsulinLevel*_ to the *peakInsulinLevel*_.
- If the BGL is greater than or equal to the *highGlucoseLevel*_, the *insulinLevel* stays at the *peakInsulinLevel*_ value.

In *CarbMetSim*, the *peakInsulinLevel*_ represents the peak ability to produce insulin. A value 1 for *peakInsulinLevel*_ means normal (or excessive, as in the case of initial stages of Type 2 Diabetes) insulin production, whereas a value 0 means that the pancreas does not produce any insulin at all (as in people with Type 1 Diabetes). A value *x* (between 0 and 1) for *peakInsulinLevel*_ means that peak insulin production is just *x* times the normal peak.

As described in the later sections, the *insulinLevel* variable has a profound impact on the operation of different organ objects in *CarbMetSim*. So, its value should be interpreted in terms of the impact it has on various organ objects, rather than the actual insulin concentration it corresponds to for a particular person. So, it is entirely possible that two very different actual insulin concentrations for two individuals map to the same value for the *insulinLevel* because they have the same impact on carbohydrate metabolism related functions of the organs.

In *CarbMetSim*’s current implementation, the *insulinLevel* variable is tightly coupled with the BGL in the manner described above. A future implementation will allow the *insulinLevel* to vary in a configurable manner by allowing the user to specify the dosage of externally injected short term insulin. This would allow the simulator to be useful for diabetes patients dependent on short term insulin injections or insulin pumps.

### 3.3 Modeling Glucose Transport

Glucose crosses cell membranes using either *active* transporters or *passive* ones. The active transporters, such as *Sodium GLucose coTransporters* (SGLTs) are able to move glucose from a low concentration to a high concentration. The passive transporters, such as *Glucose Transporters* (GLUTs) move glucose from a high concentration to a low concentration. *CarbMetSim* models the operation of active transporters in an organ by specifying the average amount of glucose transferred per minute via active transport. The actual amount transferred is a poisson distributed random variable. The simulator uses *Michaelis Menten* kinetics to determine the amount of glucose transferred in a minute via passive transport. As per the Michaelis Menten kinetics, the rate of transport (*V*) across a membrane depends on the difference in the substrate concentration (*Y*) across the membrane in the following manner: 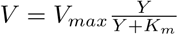, where *V*_*max*_ is the maximum rate of transport and *K*_*m*_ is the substrate concentration difference at which the transport rate is half the maximum. The *V*_*max*_ value associated with a GLUT transporter in an organ indicates the number of transporters involved. Hence, the simulator treats *V*_*max*_ associated with a particular GLUT in a particular organ as a poisson distributed random variable with a configurable mean.

#### 3.3.1 Modeling GLUT4 Operation in Muscles

Among the GLUTs, the GLUT4 transporters are of particular importance because they allow the muscles to absorb glucose from the bloodstream. When the human body is engaged in exercise, the physical activity itself *activates* sufficient number of GLUT4 transporters [42–44] and the muscles are able to absorb the desired amount of glucose from the bloodstream. *CarbMetSim* replicates this behavior. However, in the resting state, the number of *active* GLUT4 transporters depends on the insulin level in the bloodstream. When the insulin level is low (because of low BGL), GLUT4 transporters are inactive and the muscles do not absorb much glucose from the bloodstream. As the insulin level rises in the blood (in response to increase in BGL), GLUT4 transporters become active proportionately and allow the muscles to quickly absorb excess glucose from the blood.

In *CarbMetSim*, GLUT4 activation during the resting states is modeled by manipulating the *V*_*max*_ value associated with GLUT4 transporters in the following manner:

- Since a large fraction of the absorbed glucose is converted to glycogen inside muscles and there is a limit on how much glycogen can be stored inside muscles, the current amount of muscle glycogen impacts the *V*_*max*_ value. Specifically, as the muscle glycogen storage increases from zero to a configurable maximum value, the *V*_*max*_ value reduces linearly from a configurable maximum (7 mg/kg/min by default) to a configurable minimum (3.5 mg/kg/min by default).
- The impact of insulin level is captured by multiplying the *V*_*max*_ value with a factor (between 0 and 1) that increases in value with increase in the *insulinLevel*. Currently, the *insulinLevel* itself is used as the value of this factor. Since vigorous physical exercise causes temporary increase in glucose absorption by muscles [79] to make up for the glycogen lost during exercise), the *insulinLevel* does not impact the *V*_*max*_ value in the first hour after an intense physical exercise activity (unless the current BGL drops below the *baseGlucoseLevel*_).
- The impact of insulin resistance in reducing the activation of GLUT4 transporters is modeled by multiplying the the *V*_*max*_ value with a configurable parameter (*glut4Impact*_) that assumes values between 0 and 1 (by default 1.0).

### 3.4 Modeling Glycolysis

Glucose serves as a key source of energy for various tissues, which either oxidize it completely or consume it anaerobically via *glycolysis*. Complete oxidation of glucose yields 15 times more energy than anaerobic glycolysis but can only be done if oxygen is available. Tissues with access to plenty of oxygen oxidize glucose for their energy needs whereas others (possibly in the same organ) use glycolysis. Glycolysis results in the generation of lactate, which serves as a key substrate for endogenous glucose production via *gluconeogenesis* (described later).

The following organs in *CarbMetSim* use anaerobic glycolysis as an energy source: *Muscles, Liver, Kidneys, Intestine* and *Blood*. The amount of glucose consumed for glycolysis increases with the glucose availability, which is signaled by the insulin level in the bloodstream. This is modeled in the simulator in the following manner. Each organ using glycolysis as an energy source has two configurable parameters: *glycolysisMin*_ and *glycolysisMax*_ (in units of mg of glucose consumed per kg of body weight per minute). At each tick, the organ generates a poisson distributed random number (*min*) with *glycolysisMin*_ as the mean value and *glycolysisMax*_ as the maximum value. Then, subject to the glucose availability in the organ, the amount of glucose consumed in a tick for glycolysis is given by: *min* + *insulinImpact* × (*glycolysisMax*_ − *min*). Here, *insulinImpact* is a factor (between 0 and 1) that increases in value with increase in the *insulinLevel*. This factor is calculated using a sigmoid function, which is currently the CDF of a normal distribution with a configurable mean and standard deviation. The simulator also uses configurable multiplicative parameters *glycolysisMinImpact*_ and *glycolysisMaxImpact*_ (with default values 1.0) to modify the values of *glycolysisMin*_ and *glycolysisMax*_ parameters associated with each organ. These parameters can be used to model the impact of diabetes on glycolysis flux. A fraction (by default 1) of the glucose consumed for glycolysis is converted to lactate, which is added to the *Blood* object. Table 1 shows the default values for glycolysis related parameters for different organs. Here, the relative contributions of different organs towards overall glycolysis flux were set as suggested in [52, 53]. The default values of various configurable parameters in *CarbMetSim* were determined experimentally to provide a close match with published measurements performed on non-diabetic human subjects before and after a meal event [51].

**Table 1.** The default values for glycolysis related parameters in various organs.

| Organ | glycolysisMin_<br>(mg/kg/minute) | glycolysisMax_<br>(mg/kg/minute) |
| --- | --- | --- |
| Blood | 0.0315 | 0.1135 |
| Kidneys | 0.0315 | 0.1135 |
| Liver | 0.0630 | 0.5675 |
| Muscles | 0.0630 | 0.8512 |
| Intestine | 0.0315 | 0.1135 |

### 3.5 Modeling Gluconeogenesis

*Gluconeogenesis* is a metabolic pathway that allows the liver and kidneys to produce glucose from lactate, glycerol, glutamine and alanine [40, 48]. This pathway assumes special significance as the only source of glucose when no new glucose is arriving in the body via food and the glycogen store in the liver has been exhausted.

Normally, gluconeogenesis occurs when the insulin level is low (i.e. in the *post-absorptive* state). However, diabetic people may experience high gluconeogenesis flux even in the *post-prandial* state when the insulin level is high [49, 50].

In *CarbMetSim*, the *Liver* and the *Kidneys* produce glucose via gluconeogenesis at configurable average rates (*gngLiver*_ and *gngKidneys*_ respectively, 0.16 mg/kg/minute each by default) using substrates mentioned above. When the *insulinLevel* is above the *baseInsulinLevel*_ (i.e. the BGL is more than the *baseGlucoseLevel*_), the average gluconeogenesis flux is multiplied by a factor (between 0 and 1) that decreases in value with increase in the *insulinLevel* as per an inverse sigmoid function (currently, the complementary CDF of a normal distribution with a configurable mean and standard deviation). This allows us to model the decrease in gluconeogenesis flux with increase in the insulin level. On the other hand, if the *insulinLevel* is below the *baseInsulinLevel*_ (i.e. the BGL is below the *baseGlucoseLevel*_), the average gluconeogenesis flux is multiplied by a factor that decreases in value from a configurable maximum (*gngImpact*_ ≥ 1, by default 6.0) to the minimum value 1 as the *insulinLevel* increases from zero to the *baseInsulinLevel*_. This allows us to model the increased gluconeogenesis flux when BGL is low and gluconeogenesis is probably the only source of glucose for the body. The simulator currently assumes that the substrates are always available in sufficient quantity to allow gluconeogenesis to take place in the manner described above.

### 3.6 Modeling Liver Glycogen Synthesis & Breakdown

In human body, the liver helps maintain glucose homeostasis by storing excess glucose in blood during the post-prandial state (when the insulin levels are high) as glycogen and releasing glucose to the blood during the post-absorptive and exercising states (when insulin level is low) by breaking down the stored glycogen. Diabetes may effect both glycogen synthesis and breakdown in the liver.

Subject to the availability of glucose, the amount of glycogen synthesized by the *Liver* object in *CarbMetSim* simulator during each tick is a poisson distributed random variable with a configurable average (*glucoseToGlycogenInLiver*_, 4.5 mg/kg/min by default) that is modified multiplicatively by two factors. The first factor models the impact of insulin on glycogen synthesis. This factor (with values between 0 and 1) increases in value with increase in the *insulinLevel* and is calculated using a sigmoid function, which is currently the CDF of a normal distribution with a configurable mean and standard deviation. The second factor called *liverGlycogenSynthesisImpact*_ (by default 1.0) simply modifies the configured average multiplicatively and can be used to model the impact of diabetes on glycogen synthesis in the liver. The *Liver* object has a finite capacity to store glycogen and hence any excess glycogen is converted to fat and stored in the *AdiposeTissue* object.

Glycogen breakdown in the liver serves as the key source of glucose when no new glucose is entering the body via food or when the glucose needs of the body increase due to intense physical exercise. Accordingly, in *CarbMetSim*, the amount of glycogen stored in the *Liver* that is broken down to glucose during a tick closely depends on the *insulinLevel* (and hence on the current BGL). When the *insulinLevel* is above the *baseInsulinLevel*_ (i.e. the BGL is more than the *baseGlucoseLevel*_), the average glycogen breakdown flux in the *Liver* (*glycogenToGlucoseInLiver*_, 0.9 mg/kg/min by default) is multiplied by a factor (between 0 and 1) that decreases in value with increase in the *insulinLevel* as per an inverse sigmoid function (currently, the complementary CDF of a normal distribution with a configurable mean and standard deviation). This allows us to model the decrease in liver glycogen breakdown with increase in the insulin level. On the other hand, if the *insulinLevel* is below the *baseInsulinLevel*_ (i.e. the BGL is below the *baseGlucoseLevel*_), the average *Liver* glycogen breakdown flux is multiplied by a factor that decreases in value from a configurable maximum (*liverGlycogenBreakdownImpact*_ ≥ 1, by default 6.0) to the minimum value 1 as the *insulinLevel* increases from zero to the *baseInsulinLevel*_. This allows us to model the increased liver glycogen breakdown when BGL is low.

## 4 *CarbMetSim* Design and Implementation

*CarbMetSim* is a *discrete event* simulator implemented in an *object-oriented* manner. At the top level, *CarbMetSim* consists of a *SimCtl* (SIMulation ConTroLler) object and a *HumanBody* object. The *SimCtl* object maintains the simulation time (in *tick* s, where each tick is a minute) and contains a priority queue of food/exercise events sorted in order of their firing times. At the beginning of the simulation, the *SimCtl* object reads all the food/exercise events into the priority queue. At each tick, the *SimCtl* object fires the events whose firing time has arrived (by invoking appropriate methods on the *HumanBody* object) and then causes each organ to do its work during that tick (again by invoking a *HumanBody* object method).

In the following, we describe the implementation and operation of various *objects* that together implement the *CarbMetSim* simulator. The default values of various parameters listed here were determined experimentally to provide a close match with published measurements performed on non-diabetic human subjects before and after a meal event [51]. Validation of simulation results against these and other published measurements is described in later sections. Table 1 shows the default values for glycolysis related parameters for different organs. Default values of configurable parameters that determine the impact of *insulinLevel* on various metabolic processes are shown in Table 2.

**Table 2.** Configurable parameters (and their default values) for the mean and standard deviation of normal distributions to determine the impact of *insulinLevel* on various metabolic processes.

| Parameter | Default Value |
| --- | --- |
| insulinImpactOnGlycolysis_Mean | 0.5 |
| insulinImpactOnGlycolysis_StdDev | 0.2 |
| insulinImpactOnGNG_Mean | 0.5 |
| insulinImpactOnGNG_StdDev | 0.2 |
| insulinImpactGlycogenBreakdownInLiver_Mean | 0.1 |
| insulinImpactGlycogenBreakdownInLiver_StdDev | 0.02 |
| insulinImpactGlycogenSynthesisInLiver_Mean | 0.5 |
| insulinImpactGlycogenSynthesisInLiver_StdDev | 0.2 |

### 4.1 HumanBody

The *HumanBody* object serves as the container for following organ objects: *Stomach, Intestine, PortalVein, Liver, Kidneys, Muscles, AdiposeTissue, Brain, Heart* and *Blood*. At the beginning of a simulation, the *HumanBody* object does the following:

- It reads the description of various foods: their composition in terms of *rapidly*/*slowly available glucose* (RAG/SAG), protein and fat.
- It reads the description of various exercise activities: their intensity in units of *MET*s.
- It estimates the maximal rate of glucose consumption (*V O*_2_*max*, see Section 4.8) associated with the individual being simulated using the tables in Kaminsky et al. [54] given this individual’s gender, age and self-assessed fitness level within his/her age group, which are all supplied as simulation parameters.
- It reads other simulation parameters that affect the operation of different organs.

The *HumanBody* contains methods that cause the food to be added to the stomach when *SimCtl* fires a food event and update the energy needs of the body when *SimCtl* fires an exercise event. When an exercise event gets over, the *HumanBody* resets the energy needs to the resting state. When the *Stomach* has no food left, it informs the *HumanBody* about the situation. Thus, at any given time, the *HumanBody* remembers whether the stomach has some undigested food (*Fed*) or not (*PostAbsorptive*) and whether the body is currently engaged in some exercise (*Exercising*) or not (*Resting*). Accordingly, there are four body states: *Fed Resting, Fed Exercising, PostAbsorptive Resting* and *PostAbsorptive Exercising*. Different body states allow different values to be in effect for the configurable parameters governing the operation of the organs.

As mentioned before, the *HumanBody* object provides a method, which is invoked by the *SimCtl* object at each tick and causes methods to be invoked on individual organ objects that allow the organs to do their work during that tick.

### 4.2 Blood

The *Blood* object represents the bloodstream and interacts with various organs to exchange glucose, amino acids and other substrates. The *Blood* object maintains the following substrate variables: *glucose, lactate, branchedAminoAcids* (consumed by muscles, adipose tissue and brain) and *unbranchedAminoAcids*. The *Blood* object also maintains the *insulinLevel* variable discussed earlier and a *fluidVolume* variable representing the blood volume (5 liters by default). Hormones other than insulin are not currently maintained. At each tick, the *Blood* object updates the *insulinLevel* in the manner described in Seection 3.2. Also, some glucose is consumed for glycolysis in the manner described in Section 3.4.

### 4.3 Stomach

The gradual emptying of stomach contents into the intestine, also known as *gastric emptying*, is a complex phenomenon affected by a number of factors such as the volume, particle size, viscosity, osmolarity, acidity and nutritional contents of the meal [55–57]. A variety of models have been suggested in the past for the emptying of food from the stomach into the intestine. Many of these models were based on mathematical functions such as exponential [58, 59] and power exponential [60]. Lehmann and Deutsch [36] presented a simple model for gastric emptying of carboydrates in a meal, where the rate of gastric empyting has three phases - a linear increase phase, a constant maximum rate phase and a linear decrease phase. Dalla Man et al. [61] presented a three-compartment model of the gastrointestinal tract where the gastric emptying rate follows a trough-shaped pattern (initially high followed by a non-linear decrease to a minimum value followed by a non-linear increase back to the initial maximum value).

In *CarbMetSim*, when a food event is fired, the eaten food enters the *Stomach* instantaneously, where its contents are added to any existing stores of RAG, SAG, protein and fat. The simulator currently uses a simple model for gastric emptying where all the food in the stomach is assumed to be in the *chyme* form and the amount of chyme leaking to the intestine each minute consists of one part determined using a poisson distribution (with default mean 500 mg) and another part proportional to the total amount of chyme currently present in the stomach. This proportionality constant increases linearly with decrease in the energy density of the chyme. The minimum value of this proportionality constant (0.03 by default) represents the fraction leaking out of stomach each minute when the chyme consists entirely of fat (with energy density 9.0 kcal/g). On the other hand, the maximum value (9.0/4.0 times the minimum value) represents the fraction leaking out of stomach each minute when the chyme consists entirely of carbs (with energy density 4.0 kcal/g). The nutritional composition of leaked chyme is same as that of chyme present in the stomach. This simple model, inspired from [62], allows us to take in account the fat/protein induced slowdown of gastric emptying. There are many other factors that affect the gastric emptying process (the solid/liquid nature of food, fiber content, osmolarity, viscosity etc.) which *CarbMetSim* currently does not take in account. Thus, a bolus of chyme leaks from the *Stomach* into the *Intestine* every tick (i.e. every minute) until the *Stomach* is empty.

### 4.4 Intestine

#### Carbohydrate Digestion

The intestine digests the carbohydrate in the chyme using a number of enzymes to produce monosaccharides such as glucose, fructose and galactose [40]. Currently, the *Intestine* object in *CarbMetSim* converts all the carbohydrate in the chyme to just one monosaccharide - glucose. The *Intestine* receives a bolus of chyme from the *Stomach* every tick as long as there is some food in the *Stomach*. The *Intestine* maintains a list of *Chyme* objects where each object contains the undigested RAG/SAG contents of each bolus received from the *Stomach* and the time when the bolus was received. At each tick, the *Intestine* digests some amount of RAG/SAG from each *Chyme* object. The amount digested from a particular *Chyme* object is determined using normal distributions (default mean & standard-deviation: 2 minutes & 0.5 minutes for RAG and 30 minutes & 10 minutes for SAG) such that most of the RAG and SAG contents of a bolus are digested within 20 and 120 minutes respectively after the bolus’s entry into the *Intestine*. The glucose resulting from digested RAG/SAG is added to the *glucoseInLumen* variable in *Intestine*, which represents the total glucose present in the intestinal lumen. This glucose is processed as described later in this section.

#### Fat and Protein Digestion

As a chyme bolus enters the *Intestine* from the *Stomach*, its fat contents are simply added to the *AdiposeTissue* and its protein contents are added to a common protein pool in *Intestine*. At each tick, the *Intestine* digests a small amount of this protein (determined as per a poisson distribution with default mean 1mg) and transfers the resulting amino acids to the *PortalVein*. The simulator does not keep track of the amino acid contents of dietary protein and makes a simple assumption that 85% of these amino acids are *unbranched* and the remaining 15% are *branched*.

#### Glucose Absorption from Intestine to PortalVein

The glucose moves from the intestinal lumen to the enterocytes across the brush border membrane and then from the enterocytes to the portal vein across the basolateral membrane. The transfer from the intestinal lumen to the enterocytes takes place via a combination of active (SGLT-1) and passive (GLUT2) transporters, where the number of GLUT2 transporters in action depends on the glucose concentration on the lumen side. The transfer from the enterocytes to the portal vein takes place solely via passive GLUT2 transporters [40]. The *Intestine* object maintains two variables: *glucoseInLumen* and *glucoseInEnterocytes*, which represent total glucose present in the intestinal lumen and in enterocytes respectively. At each tick, the *Intestine* moves some glucose from *glucoseInLumen* to *glucoseInEnterocytes*. The amount moved has an active transport component (poisson distributed with default mean 30 mg/minute) and a passive transport component determined using Michaelis Menten kinetics (assuming configurable volumes for the lumen and the enterocytes). The *V*_*max*_ value used for Michaelis Menten kinetics increases with glucose concentration in the lumen with default maximum value 800 mg/minute. The *K*_*m*_ value used is 20 mmol/l by default [40]. Glucose transport from the enterocytes to the portal vein is modeled by moving some glucose from *glucoseInEnterocytes* to the *PortalVein* at each tick. The amount moved is determined using Michaelis Menten kinetics (average *V*_*max*_ = 800 mg/minute, *K*_*m*_ = 20 mmol/l by default [40]).

#### Glycolysis

The intestinal cells get some of their energy via glycolysis of glucose to lactate in the manner described in Section 3.4. If the glucose in enterocytes (*glucoseInEnterocytes*) is not sufficient, the extra glucose needed for glycolysis comes from the bloodstream (the *Blood* object).

### 4.5 PortalVein

The portal vein carries blood that has passed through the intestinal tract to the liver. Due to its special status as the conduit from the intestine to the liver, *CarbMetSim* maintains the portal vein as a separate entity (the *PortalVein* object) from rest of the circulatory system (represented by the *Blood* object). The glucose and amino acids resulting from the food digestion in the *Intestine* travel to the *Liver* via the *PortalVein*.

Since the portal vein is a part of the circulatory system, it must have the same glucose concentration as rest of the circulatory system when no new glucose is being received from the intestine. This is achieved in *CarbMetSim* in the following manner. At the beginning of a tick, there is no glucose in the *PortalVein*. During each tick, the following sequence of actions take place:

- The *PortalVein* imports glucose from the *Blood* so that the glucose concentration in the *PortalVein* matches that of *Blood* before the import. The *PortalVein*’s volume, used to calculate the glucose concentration, is a configurable parameter with default value 5 dl.
- Glucose transfer takes place from the *Intestine* to the *PortalVein* (as described previously in Section 4.4) and then from the *PortalVein* to the *Liver* (as described next in Section 4.6).
- Finally, any remaining glucose in the *PortalVein* is moved back to the *Blood*.

All the amino acids received from the *Intestine* into the *PortalVein* during a tick are moved to the *Liver* during that tick itself.

### 4.6 Liver

The *hepatocytes* in the liver absorb glucose from the portal vein via GLUT2s when the glucose concentration in the portal vein is higher. The absorbed glucose is phosphorylated to glucose 6-phosphate, which is used either for glycogen synthesis or for glycolysis. Insulin and glucose activate the enzymes associated with glycogen synthesis and inhibit those associated with glycogen breakdown. Insulin also activates glycolysis of glucose 6-phosphate in hepatocytes to form pyruvate, some of which is oxidized and the remaining is converted to lactate and released to the bloodstream. On the other hand, lack of insulin (and presence of glucagon) activates glycogen breakdown (to glucose) as well as gluconeogenesis (which again produces glucose). The gluconeogenesis flux increases with the availability of the substrates (such as lactate, alanine and glycerol) in the bloodstream even if the insulin level is high. Excess glucose in the hepatocytes is either used for glycogen synthesis (if the insulin level is high) or leaves the cells via GLUT2s and possibly other means (if the insulin level is low). High insulin level also causes some of the excess glucose to be converted to lipid. Thus, the liver absorbs glucose during the fed state and uses it for glycogen synthesis and glycolysis. On the other hand, the liver releases glucose to the bloodstream during the post-absorptive and exercising states via glycogen breakdown and gluconeogenesis. Another important aspect of the liver operation is its oxidation of *unbranched* amino acids which provides for almost half of the liver’s energy requirements.

*CarbMetSim* implements the liver operation in the *Liver* object. The simulator allows the initial amount of glycogen stored in the *Liver* as well as the maximum amount it can hold to be set via configurable parameters. By default, the *Liver* has sufficient glycogen at the beginning of a simulation to produce 100 grams of glucose. Also, by default, an amount equivalent to 120 grams of glucose is the upper limit on the amount of glycogen that the *Liver* object can store. At each tick, the *Liver* does the following:

- *Glucose Absorption*/*Release:* If the glucose concentration in higher in the *PortalVein* than in the *Liver*, some glucose will be absorbed in the *Liver* via GLUT2s. Similarly, if the glucose concentration is higher in the *Liver* than in the *Blood*, some glucose will be released to the *Blood* via GLUT2s. The amount of the glucose absorbed/released is determined using Michaelis Menten kinetics (with default average *V*_*max*_=50mg/kg/min and default *K*_*m*_=20 mmol/l [40]).
- *Glycogen Synthesis*/*Breakdown:* The *Liver* performs glycogen synthesis or breakdown in the manner described in Section 3.6.
- *Lipogenesis:* If the glycogen storage in the *Liver* exceeds its maximum configured value, the excess glycogen is converted to fat, which is stored in *AdiposeTissue*.
- *Glycolysis* and *Gluconeogenesis:* The *Liver* consumes some glucose for glycolysis in the manner described in Section 3.4 and produces glucose via gluconeogenesis in the manner described in Section 3.5.
- *Amino Acid Consumption:* The *Liver* consumes 93% of unbranched amino acids received from the *PortalVein* and releases the rest (along with all the branched amino acids) to the *Blood* object.

### 4.7 Kidneys

The kidneys filter the blood and require significant amount of energy for this task. Their outer layer (the *cortex*) is well supplied with oxygen and hence meets its energy needs via oxidation of glucose and fatty acids absorbed from the bloodstream. The inner core (the *medulla*) uses anaerobic glycolysis for energy. The kidneys also generate glucose via gluconeogenesis. *CarbMetSim* implements the kidney operation in the *Kidneys* object. At each tick, the *Kidneys* do the following:

- *Glycolysis:* The renal medulla in *Kidneys* meets its energy requirements via glycolysis, which is implemented in the manner described in Section 3.4. The glucose consumed for glycolysis is absorbed from the *Blood* object and the resulting lactate is released to the *Blood* object.
- *Gluconeogenesis:* The *Kidneys* produce glucose via gluconeogenesis in the manner described in Section 3.5 and release it to the *Blood* object.
- *Glucose Excretion in Urine:* As the the glucose concentration in *Blood* increases from one threshold (11 mmol/l [52, 64] by default) to another (22 mmol/l by default), the glucose excretion in urine increases linearly from zero to a certain peak level (100 mg/min by default). The simulator supports a configurable parameter *excretionKidneysImpact*_ (with default value 1) to multiplicatively modify the amount of glucose excreted per tick in urine.

### 4.8 Muscles

The skeletal muscles have two types of cells or *fibers*: the *red* fibers oxidize substrates (fatty acids, glucose) absorbed from the bloodstream to meet their energy needs while the *white* fibers rely on glycolysis of glucose 6-phosphate obtained from the glycogen stored within the white fibers for energy. The glucose absorption from the bloodstream occurs mainly via insulin-sensitive GLUT4 transporters with some *basal* level absorption taking place via GLUT1 transporters. The skeletal muscles also use some *branched chain* amino acids absorbed from the bloodstream to meet their energy needs.

#### Muscles Operation During Rest [40, 47, 65]

In the resting state, the muscles meet 85 − 90% of their energy needs via the oxidation of fatty acids. About 10% of the energy comes from oxidation of glucose and 1 − 2% from amino acids. The glucose is absorbed from the bloodstream using GLUT4 and GLUT1 transporters as mentioned earlier. The absorbed glucose is used for oxidation, glycogen synthesis and glycolysis [66]. The glucose oxidation and glycolysis in muscles under resting conditions increases with the insulin level in the bloodstream.

#### Muscles Operation During Aerobic Activity [40, 47, 63]

Oxidation of glucose and fatty acids is the main source of energy for exercising muscles. The relative fraction of these substrates used to meet the energy needs depends on the exercise intensity, which in turn is decided based on the rate at which the individual consumes oxygen while doing this exercise. Each individual has a certain maximal rate at which he/she can consume oxygen. This individual-specific maximal rate, called *V O*_2_*max*, depends on the gender, age and fitness level of the individual [54]. So, the intensity of an exercise activity can be described by the oxygen consumption rate (as the %age of the individual’s *V O*_2_*max*) associated with this exercise. The exercise intensity can also be described in an individual-independent manner in units of *Metabolic Equivalent of Task* or *MET*s, where 1 *MET* is 1 *kcal* of energy expenditure per kg of body weight per hour. By convention, 1 *MET* is considered equivalent to 3.5*ml* of oxygen consumption per kg of body weight per minute. So, an exercise with a certain intensity in terms of *MET*s may translate to very different intensities in terms of %*V O*_2_*max* for different individuals.

Romijn et al. [63] reported that about 10% of the energy needs during aerobic exercise are met by oxidizing glucose absorbed from the blood via GLUT4/GLUT1 transporters. The aerobic activity is sufficient to activate GLUT4 transporters. So, their action is not dependent on the insulin during the aerobic exercise [42–44]. For low intensity (e.g. 25%*V O*_2_*max*) exercise, almost all of the remaining energy needs are met by oxidizing fatty acids [63]. For moderate and high intensity exercise, a significant fraction of energy needs is met by oxidation of glucose derived from the glycogen stored locally in the exercising muscles. Romijn et al. [63] reported about 30% of the energy coming from the oxidation of glucose derived from locally stored glycogen when the intensity of the aerobic exercise was 65%*V O*_2_*max*. Horton [47] reported oxidation of glucose (absorbed from the blood and derived from local glycogen) providing for about 50% and almost 100% of the energy needs when the exercise intensities were 50%*V O*_2_*max* and 100%*V O*_2_*max* respectively. Most of the remaining energy needs are met by oxidation of fatty acids [67]. Once the glycogen stored in the liver and the muscles is over, it becomes impossible for the individual to perform very high intensity exercise. A small fraction of the energy needs is met by glycolysis of glucose 6-phosphate derived from locally stored glycogen. The glycolysis level increases linearly with exercise intensity. Finally, a very small fraction of energy needs is met by consuming *branched* amino acids absorbed from the blood [67].

#### 4.8.1 Implementation in *CarbMetSim*

In *CarbMetSim*, the skeletal muscles are implemented as the *Muscles* object. Currently, the simulator implements response to the resting condition and the aerobic exercise only. Specifically, exercise with a significant anaerobic component cannot yet be simulated. Also, it is not yet possible to distinguish among different muscle groups.

At the beginning of a simulation, the *HumanBody* object estimates the *V O*_2_*max* associated with the individual being simulated using the tables in Kaminsky et al. [54] given this individual’s gender, age and self-assessed fitness level within his/her age group, which are all supplied to the simulator as input parameters. When an exercise event is fired, the exercise intensity is translated from the units of *MET*s into %*V O*_2_*max*. The exercise intensity determines the fraction of the energy needs met via oxidation of glucose derived from locally stored glycogen. The simulator allows the initial amount of glycogen stored in the *Muscles* as well as the maximum amount it can hold to be set via configurable parameters. By default, both these parameters have values equivalent to 500 grams of glucose.

When the *HumanBody* is in *Fed*_*Exercising* or *PostAbsorptive*_*Exercising* state during a tick, the *Muscles* object performs the following actions:

- *Oxidation of glucose absorbed from the* Blood: The *Muscles* absorb a random amount of glucose from the *Blood* (up to a configurable limit which is 30*μ*mol/kg/min by default) so that it can be oxidized to meet on average 10% of the energy needs during this tick. This absorption does not depend on the current *insulinLevel* in the *Blood*.
- *Oxidation of glucose derived from local glycogen:* The exercise intensity (in %*V O*_2_*max*) is used to determine the fraction of energy needs that will be met by oxidizing glucose derived from locally stored glycogen. As the exercise intensity increases from 0%*V O*_2_*max* to 100%*V O*_2_*max*, a value between 0 and 0.9 is determined using a sigmoid function (currently, the compressed CDF of a normal distribution) such that exercise intensities 50%*V O*_2_*max* and 100%*V O*_2_*max* yield values close to 0.4 and 0.9 respectively. This value is then used as the mean to generate a random value that gives the fraction of energy needs during this tick to be met by oxidizing glucose derived from local glycogen (as long as a sufficient amount of local glycogen is available).
- *Glycolysis:* The glycolysis flux during the tick increases linearly from a (poisson distributed) random value (with *glycolysisMin*_ as the mean) to *glycolysisMax*_ as the exercise intensity increases from 0%*V O*_2_*max* to 100%*V O*_2_*max*. The glucose 6-phosphate consumed for glycolysis comes from locally stored glycogen. The resulting lactate is added to the *Blood* object.
- *Fatty Acid Consumption:* If glucose oxidation and glycolysis described above do not meet the current energy needs, *Muscles* consume fat (representing fatty acids) from the *AdiposeTissue* to meet the remaining energy needs.

When the *HumanBody* is in *Fed*_*Resting* or *PostAbsorptive*_*Resting* state during a tick, the *Muscles* object performs the following actions:

- *Glucose Absorption:* GLUT4 based glucose absorption [40] occurs in the manner described in Section 3.3.1. Also, basal absorption via GLUT1s occurs at a configured rate (by default zero).
- *Glycolysis:* A fraction of the absorbed glucose (determined as described in Section 3.4) is consumed via glycolysis and the resulting lactate is added to the *Blood* object.
- *Glycogen Synthesis:* If the glycogen store of the *Muscles* is less than the maximum amount that *Muscles* could hold [40], a (poisson distributed) random amount of the absorbed glucose (with a configurable mean, 7.0mg/kg/min by default) is converted to glycogen.
- *Oxidation:* Remainder of the absorbed glucose is considered consumed via oxidation.
- *Fatty Acid Consumption:* If glycolysis and glucose oxidation described above do not meet the energy needs during the resting state, *Muscles* consume fat (representing fatty acids) from the *AdiposeTissue* to meet the remaining energy needs.

### 4.9 Adipose Tissue

*CarbMetSim* does not yet have a detailed implementation of lipid metabolism. Currently, the *AdiposeTissue* object serves as the storage for fat. The *Intestine* object directly adds the fat contents in chyme to the *AdiposeTissue* object. Similarly, the *Liver* object converts excess glycogen to fat to be stored in the *AdiposeTissue* object. The *Muscles* object directly removes fat from the *AdiposeTissue* in accordance with its energy needs.

### 4.10 Brain

The brain meets its energy needs by oxidizing glucose (although under starvation conditions it can also use ketone bodies). The *nerve* cells in the brain use GLUT3 transporters to absorb glucose from the bloodstream. Since the *K*_*m*_ value associated with GLUT3 transporters is quite low, the rate of glucose absorption by the nerve cells does not change much with glucose concentration in the bloodstream (unless it drops way below the normal levels). The brain oxidizes about 120 g of glucose per day, equivalent to absorption of about 83.33 mg of glucose per minute [40, 66]. In *CarbMetSim*, the brain operation is modeled as the *Brain* object which consumes a (poisson distributed) random amount (with mean 83.33 mg) of glucose every minute from the *Blood* object.

### 4.11 Heart

The heart meets most of its energy needs by oxidizing fatty acids. Depending upon their availability, up to 30% of the heart’s energy needs are met by consuming glucose and lactate [68]. A much smaller part of the energy needs is met from amino acids and ketone bodies. The heart uses both GLUT1 and GLUT4 transporters to absorb glucose from the bloodstream. The *Heart* object in *CarbMetSim* models the heart operation. It absorbs a poisson distributed random amount of glucose (with default mean 14 mg/minute [66]) from *Blood* and oxidizes it to meet its energy needs.

## 5 Validation of *CarbMetSim* For a Meal Event

In order to determine the default values of various configurable parameters for normal subjects and to illustrate *CarbMetSim*’s ability to model carbohydrate metabolism in normal subjects and those with Type 2 Diabetes (T2D) in *post-absorptive* and *post-prandial* phases, we configured the simulator to provide a close match with measurements reported in Woerle et al. [51]. Woerle et al. [51] did extensive measurements on 26 subjects with Type 2 Diabetes and 15 age/weight/sex-matched subjects without diabetes to determine the flux along different pathways for glucose arrival and consumption following a standard meal. The T2D subjects included 16 men and 10 women with following characteristics: age 53 ± 2 years, body weight 93 ± 4 kg, BMI 30 ± 1*kg*/*m*^2^, body fat 34 ± 3 %, average *HbA1c* 8.6 ± 0.3%. The normal subjects included 7 men and 8 women with following characteristics: age 49 ± 3 years, body weight 89 ± 4 kg, BMI 30 ± 1*kg*/*m*^2^, body fat 36 ± 3 %. All the subjects consumed a standard breakfast, consisting of 84 g of glucose, 10 g of fat and 26 g of protein, at 10am on the day of the measurements after a fast of more than 14 hours. Measurements were performed for the post-absorptive phase before the breakfast and the post-prandial phase assumed to be six hours in duration after the breakfast.

Two sets of simulations were performed using *CarbMetSim*: one for a normal subject and one for a T2D subject (see Table 3). Each set consisted of 30 simulations with different seeds for random number generation. Default values were used for most of the configurable simulation parameters. These values (already reported in the previous sections) were set so that the normal subject simulations achieves a close match with measurements reported in [51]. In particular, the impact of insulin level on the gluconeogenesis flux was disabled because insulin level did not seem to influence the gluconeogenesis flux in the reported measurements. Configurable parameters for which the default values were not used are shown in Table 3. In all the simulations reported in this section and the remaining ones, the parameters did not change in value with the body state (although the simulator is capable of using different values for a parameter depending on the body state). Simulation parameters *bodyWeight* and *age*_ were set to the average values reported for subjects in each category in [51]. Parameters *age*_, *gender*_ and *fitnessLevel*_ are used to determine *V O*_2_*max*, the maximal rate of oxygen consumption for the subject being simulated and are not relevant for the simulations reported in this section. The following parameters were set as per the data reported in [51] (see Tables 4 and 5):

**Table 3.** Configuration parameters for simulations for a single meal event.

|  | Normal | Type 2 Diabetic |
| --- | --- | --- |
| age_ (years) | 49 | 53 |
| gender_ | 0 (male) | 0 (male) |
| fitnessLevel_ (%ile) | 50 | 50 |
| bodyWeight (kg) | 89 | 93 |
| minGlucoseLevel_ (mg/dl) | 50 | 50 |
| baseGlucoseLevel_ (mg/dl) | 90 | 210 |
| highGlucoseLevel_ (mg/dl) | 145 | 360 |
| baseInsulinLevel_ | 0.001 | 0.001 |
| peakInsulinLevel_ | 1.0 | 0.6 |
| glut4Impact_ | 1.0 | 0.25 |
| glycolysisMinImpact_ | 1.0 | 4.0 |
| glycolysisMaxImpact_ | 1.0 | 1.5 |
| excretionKidneysImpact_ | 1.0 | 1.3 |
| glucoseToGlycogenInLiver_ (mg/kg/min) | 4.5 | 6.75 |
| glycogenToGlucoseInLiver_ (mg/kg/min) | 0.9 | 1.25 |
| gngLiver_ (mg/kg/min) | 0.16 | 0.38 |
| gngKidneys_ (mg/kg/min) | 0.16 | 0.38 |

**Table 4.** Normal Subjects: Key Measurements From Woerle Et Al. [51] and Corresponding Results From 30 Simulations with Different Seeds. “Before Breakfast” Simulation Results Were Observed at 9.59AM. All Values Expressed as *Mean* ± *Std Error*.

|  | Woerle et al. [51] | Simulations |
| --- | --- | --- |
| **Before Breakfast** |  |  |
| BGL | $4.7 \pm 0.1$ mM ($84.6 \pm 1.8$ mg/dl) | $91.937 \pm 0.010$ mg/dl |
| Glycogen Breakdown | $5.5 \pm 0.6$ μmol/kg/min ($88.1 \pm 9.6$ mg/min) | $80.064 \pm 0.171$ mg/min |
| Gluconeogenesis | $2.6 \pm 0.2$ μmol/kg/min ($41.6 \pm 3.2$ mg/min) | $41.720 \pm 0.053$ mg/min |
| **90 Minutes After Breakfast** |  |  |
| Plasma Insulin | $290 \pm 29$ pM | $0.993 \pm 0.001$ |
| Glycogen Breakdown | $1.3 \pm 0.6$ μmol/kg/min ($20.8 \pm 9.6$ mg/min) | 0 mg/min |
| Gluconeogenesis | $2.6 \pm 0.2$ μmol/kg/min ($41.6 \pm 3.2$ mg/min) | $41.824 \pm 0.051$ mg/min |
| Peak Post-prandial BGL | 8 mM (144 mg/dl) | $144.826 \pm 0.046$ mg/dl |
| **Total Glucose Consumed/Produced During 6 Hours After The Breakfast** |  |  |
| Gluconeogenesis | $15.3 \pm 1.2$ g | $15.041 \pm 0.001$ g |
| Glycogen Breakdown | $4.3 \pm 1.7$ g | $16.241 \pm 0.002$ g |
| Glucose Excretion in Urine | $0.7 \pm 0.4$ g | 0 g |
| Oxidation | $45.6 \pm 2.6$ g | $47.055 \pm 0.008$ g |
| Glycolysis | $21.5 \pm 2.2$ g | $22.614 \pm 0.002$ g |
| Glycogen Storage | $40.6 \pm 3.6$ g | $45.616 \pm 0.008$ g |

**Table 5.**
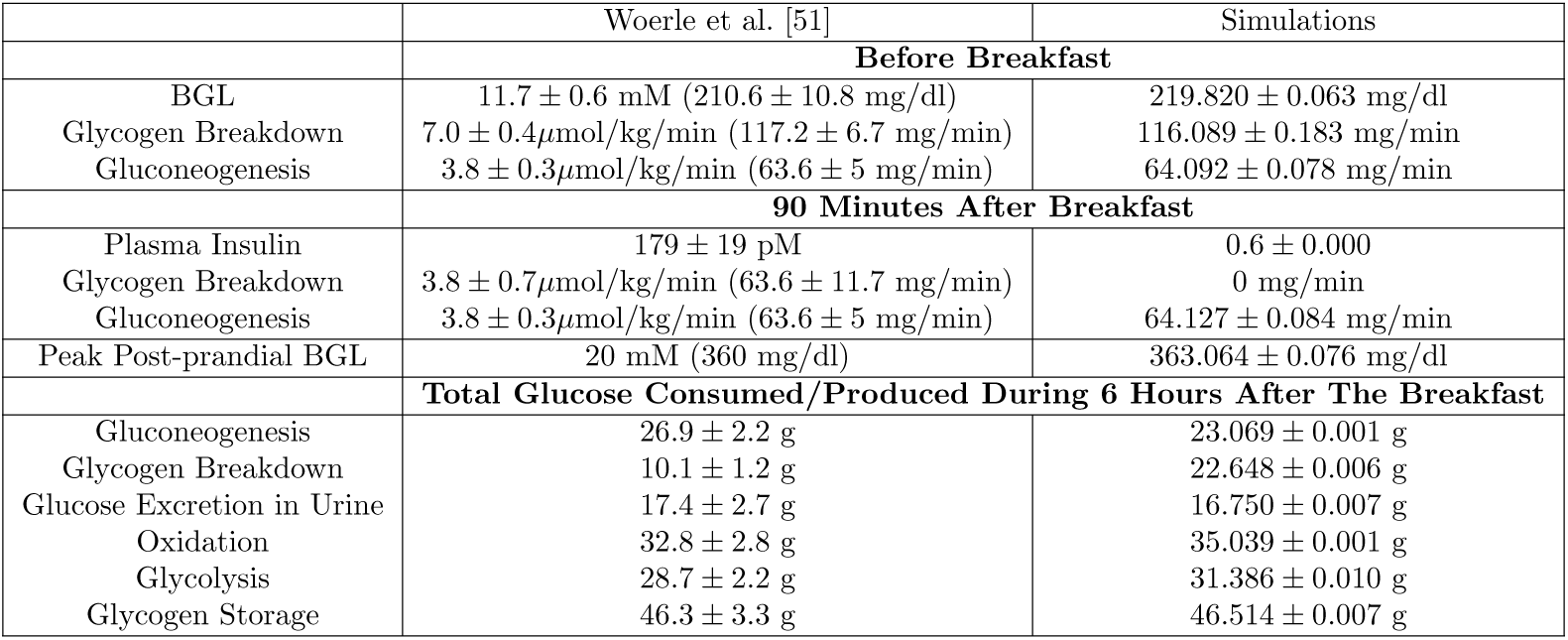
Subjects with Type 2 Diabetes: Key Measurements From Woerle Et Al. [51] and Corresponding Results From 30 Simulations with Different Seeds. “Before Breakfast” Simulation Results Were Observed at 9.59AM. All Values Expressed as *Mean* ± *Std Error*.

- The *baseGlucoseLevel*_ and *highGlucoseLevel*_ values were set to the reported values for the fasting and the peak BGL [51].
- The *peakInsulinLevel*_ values were set according to the reported peak plasma insulin levels [51].
- The average glycogen breakdown flux in the *Liver* (*glycogenToGlucoseInLiver*_) values was set to achieve a good match with the reported values for the *post-absorptive* glycogen breakdown flux and the total glycogen breakdown in the liver during the *post-prandial* phase.

Each simulation ran for 18 hours of simulated time: from 12am in midnight till 6pm in the next evening with one meal (consisting of 84 g of glucose, 10 g of fat and 26 g of protein) intake event happening at 10am. There were no other events during the simulated time and the simulated subject was already in the post-absorptive state when the simulation started at 12am.

Fig 1 shows the minute-by-minute values of interest in two simulations with a particular seed value (for random number generation): one for the normal subject and the other for the T2D subject. Fig 1a shows that the gastric emptying is complete within 45 minutes of meal intake. The fat contents of the meal was responsible for some of the delay in gastric emptying. Fig 1b shows the rapid digestion of glucose as it arrives in the *Intestine* and Fig 1c shows the appearance of digested glucose in the *PortalVein* as described in Section IV-D. Fig 1d shows the change in BGL throughout the post-prandial phase starting from the post-absorptive levels before 10am.

**Fig 1.**
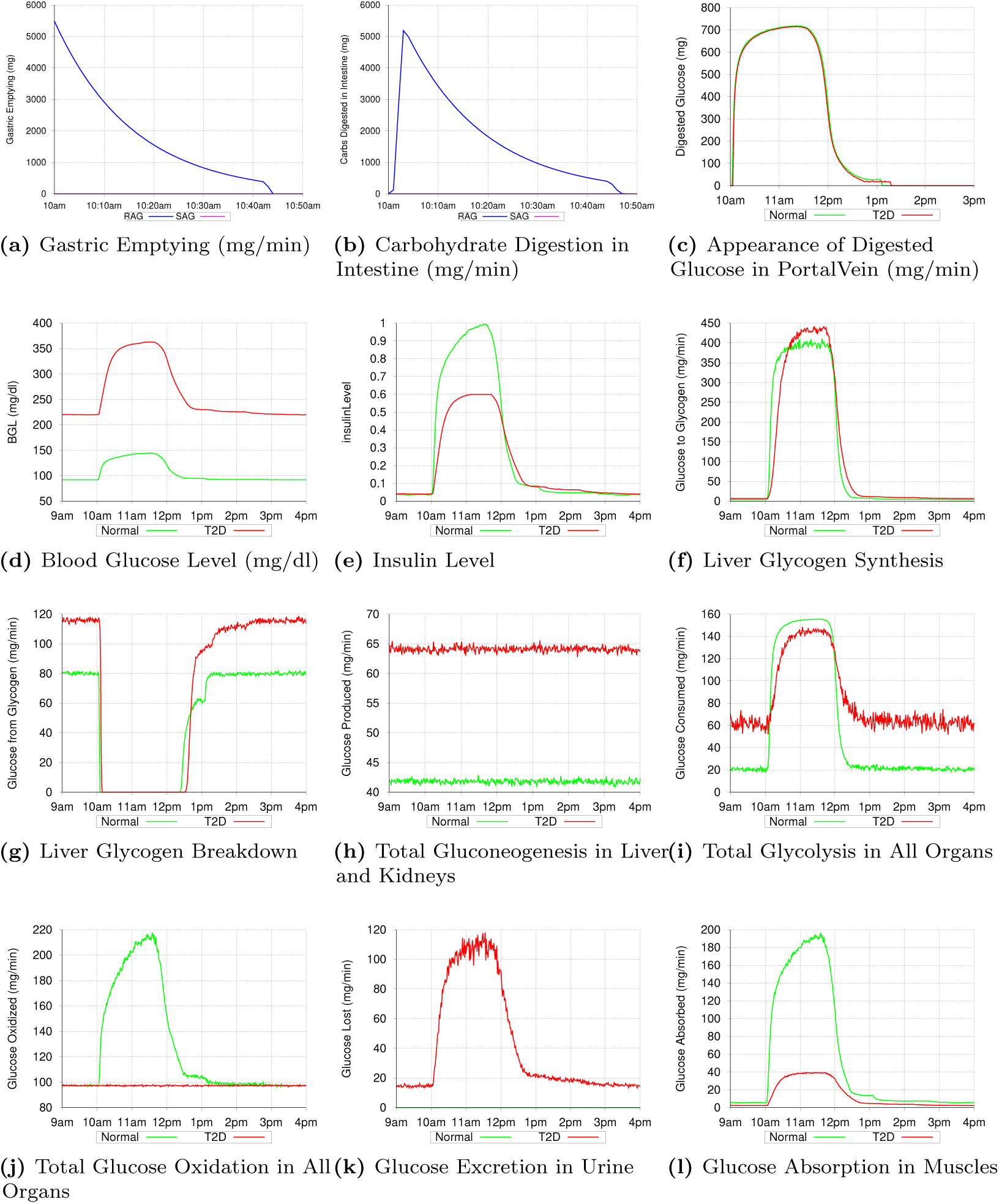
Simulating a Meal Event for a Normal Subject and a Subject with Type 2 Diabetes (T2D): Minute-by-minute Values For Important Processes In Simulations With a Particular Seed for Random Number Generation.

### Post-Absorptive Phase

During the post-absorptive phase (before 10am), the *insulinLevel* (Fig 1e) is low enough to ensure that the glucose production via glycogen breakdown in the *Liver* (Fig 1g) takes place at the peak level, there is no glycogen synthesis in the Liver (Fig 1f) and glucose consumption via oxidation (Fig 1j) & glycolysis (Fig 1i) in various organs is at their minimum levels. Gluconeogenesis (Fig 1h) takes place in the *Liver* and the *Kidneys*, unaffected by the *insulinLevel* (as reported in [51]), at the configured rates specified in [51] and provides the second source of glucose during the post-absorptive phase. While the minimum glucose oxidation flux is largely determined by the needs of the *Brain* and *Heart*, the configured values for the minimum glycolysis flux are chosen so that total glucose consumption during the post-absorptive phase matches the glucose production during this phase. Accordingly, the *glycolysisMinImpact*_ parameter was set to value 4.0 in simulations for the T2D subject and hence the glycolysis flux for the T2D subject during the post-absorptive phase is much higher than that for the normal subject (Fig 1i). Thus, during the post-absorptive phase, total glucose production (glycogen breakdown + gluconeogenesis) is matched closely by the total glucose consumption (oxidation + glycolysis + excretion in urine) and the BGL stabilizes to a value near the *baseGlucoseLevel*_. Since the glycogen breakdown in *Liver* is configured to rapidly slow down with increase in the *insulinLevel*, any temporary mismatch between glucose production and consumption is quickly corrected.

### Post-Prandial Phase

The post-prandial phase in these simulations begins with the meal intake at 10am. The BGL begins to rise (as shown in Fig 1d) with the arrival of the digested glucose in the *PortalVein*. Increase in the BGL causes the *insulinLevel* to increase (Fig 1e) which rapidly brings glycogen breakdown in the *Liver* to a halt (Fig 1g). However, the influx of digested glucose (≈ 700 mg/minute at peak) is more than sufficient to compensate for the halt in glycogen breakdown (peak value ≈ 120 and 80 mg/minute for T2D and normal subjects respectively) and the BGL (and hence the *insulinLevel*) continues to rise. Glucose production via gluconeogenesis (Fig 1h) continues as before unaffected by the increase in the *insulinLevel* (as reported in [51]). In the simulation for the normal subject, increase in the *insulinLevel* causes a proportional increase in the GLUT4 activation and hence in the glucose absorption by *Muscles* (Fig 1l). The glucose absorbed by the *Muscles* is consumed via glycolysis and oxidation. Since the glycogen stores in the *Muscles* are already full, none of the absorbed glucose is stored as glycogen in the *Muscles*. Increased oxidation flux seen in Fig 1j between 10am and 1pm in the normal subject is mainly due to the increased glucose oxidation in the *Muscles*. Impaired GLUT4 activation in the T2D subject (caused by *glut4Impact* value 0.25) means that the diabetic *Muscles* are not able to absorb as much glucose as the normal *Muscles* (Fig 1l). Also, almost all of the glucose absorbed by the diabetic *Muscles* is consumed via glycolysis (since the oxidation flux shown in Fig 1j does not show any rise in the post-prandial phase for the T2D subject). This is because of the much higher value of the minimum glycolysis flux in the T2D subject than for the normal subject (caused by the *glycolysisMinImpact*_ parameter having value 4.0 for the T2D subject and 1.0 for the normal subject). Increase in the *insulinLevel* causes glycolysis flux to increase in other organs too (Fig 1i). The peak glycolysis flux for the T2D subject is configured to be higher than that for the normal subject (by setting *glycolysisMaxImpact*_ to 1.25 for the T2D subject) so as to achieve a close match with reported results in [51] for the total glycolysis flux during the *post-prandial* phase assumed to be between 10am and 4pm (see Table 5). As the BGL approaches the *highGlucoseLevel*_ (and *insulinLevel* approaches the *peakInsulinLevel*_), glycogen synthesis in the *Liver* starts and quickly ramps up to its peak level (see Fig 1f) thereby significantly slowing down any further increase in BGL. Note that the total glycogen storage during the *post-prandial* phase as reported in [51] (and shown in Tables 4 and 5) is higher for the T2D subjects than for the normal subjects even though the T2D subjects have much smaller peak insulin levels. As described in Section 3.6, the *insulinLevel* has a big impact on glycogen synthesis in the *Liver*. In order to compensate for lower insulin levels in the T2D subjects, the *glucoseToGlycogenInLiver*_ parameter in the simulations is assigned a much higher value for the T2D subject than for the normal subject (6.75 mg/kg/min versus 4.5 mg/kg/min). For the T2D subject, a significant amount of glucose is also lost via excretion in urine (see Fig 1k). Thus, BGL stays around the *highGlucoseLevel*_ as long as the digested glucose is appearing in the *PortalVein* at the peak rate. As the digested glucose appearance in the *PortalVein* slows down, the BGL begins to drop and the glycogen synthesis in the *Liver* quickly comes to a halt thereby slowing down the rate at which the BGL falls. Decrease in BGL also causes decrease in the glycolysis flux, the glucose absorption by the *Muscles* and the glucose excretion in the urine, which further slows down the rate of BGL decrease. As BGL approaches the *baseGlucoseLevel*_, the glycogen breakdown in the *Liver* quickly ramps up to prevent any further decrease in the BGL and another post-absorptive phase begins.

### Comparison with Measurements from Woerle et al. [51]

Tables 4 and 5 show the key measurements from [51] for normal and T2D subjects respectively along with the corresponding results from the simulations. The simulations were configured to use the post-absorptive (peak) glycogen breakdown and gluconeogenesis flux values reported in [51]. With appropriate settings for other configurable parameters (Table 3), the post-absorptive BGLs in the simulations were close to the values reported in [51] for both normal and T2D subjects. The simulations were configured to ensure that the *insulinLevel* does not have any impact on gluconeogenesis flux (as reported in [51]). Accordingly, the gluconeogenesis flux in simulations 90 minutes after the breakfast was same as that before the breakfast matching the numbers reported in [51]. The glycogen breakdown 90 minutes after the breakfast was still substantial in [51] but had completely halted in the simulations. Overall, the peak post-prandial BGLs in simulations matched the ones reported in [51]. Total glucose produced/consumed along various pathways in the simulations for the normal subject during 6 hours after the breakfast was similar to values reported in [51]. The only exception was glycogen breakdown. It is clear from Fig 1 that the post-prandial phase in simulations was over by 1pm and hence glycogen breakdown happened at the peak level between 1pm and 4pm (Fig 1g). Apparently, this was not the case during measurements reported in [51] and the average glycogen breakdown flux during 6 hours after the breakfast was quite low. This combined with the fact that the glycogen breakdown was still substantial 90 minutes after the breakfast (when insulin levels were at their peak) means that glycogen breakdown process is relatively slow in reacting to the insulin levels. We observed a similar mismatch between the simulation results for the T2D subject and the values reported in [51] for total glycogen breakdown during 6 hours after the breakfast. Also, for the T2D subjects, [51] reported somewhat higher total gluconeogenesis flux during 6 hours after the breakfast (26.9 ± 2.2 g) than what we observed in the simulations (23.069 ± 0.001 g). Since gluconeogenesis flux in the simulation had same values during both post-prandial and post-absorptive phase, higher total flux reported in [51] means that gluconeogenesis flux actually increased during the post-prandial phase (perhaps due to higher availability of gluconeogenesis substrates). The simulator currently does not support increase in gluconeogenesis flux due to increased availability of substrates. Other results for T2D subjects in [51] were quite similar to what we observed in the simulations. Overall, it can be said that the simulation results closely matched those reported in [51] for both normal and T2D subjects.

## 6 Validation of *CarbMetSim* for an Exercise Event

Carbohydrate metabolism during and after an exercise event has been extensively studied for both normal and diabetic subjects [44, 47, 69–72]. As described in Section 4.8, glucose oxidation plays a major role in meeting energy needs during physical exercise. The exercising muscles get the glucose they need by breaking down locally stored glycogen and by absorbing glucose from the bloodstream. The glucose absorption from the bloodstream does not depend on the insulin levels since the physical exercise itself is sufficient to activate GLUT4 transporters [42–44].

In case of normal people, physical exercise inhibits insulin secretion [65, 73–76] and promotes secretion of other hormones such as glucogon [65, 75–77]. These changes allow the liver to break sufficient glycogen to meet the increased glucose needs. So, as long as glycogen is available in the liver and in the exercising muscles, the glucose production via glycogenolysis (in the liver and the exercising muscles) and gluconeogenesis (in the liver and kidneys) generally matches the glucose consumption by exercising muscles (and other organs) and the blood glucose level stays in the normal range [44, 47]. The blood glucose level will drop once the glycogen stores have been exhausted and gluconeogenesis alone is not sufficient to match the glucose consumption by the exercising muscles. In case of people with Type 2 Diabetes or insulin-treated Type 1 Diabetes, physical exercise fails to sufficiently reduce insulin level in the blood and as a result the glycogen breakdown in the liver may not be sufficient to meet the additional glucose needs [78]. Thus, the blood glucose level may drop significantly during physical exercise. Finally, people with Type 1 Diabetes with too little insulin in their system may experience an increase in BGL (which was already high before the exercise) when they indulge in physical exercise. This may happen because the secretion of glucagon and other hormones during exercise increases glucose production via glycogen breakdown in the liver to a rate much higher than the impaired rate at which the exercising muscles absorb glucose in some people with Type 1 Diabetes [44–47].

In order to demonstrate *CarbMetSim*’s ability to simulate the impact of aerobic physical exercise, we report in this section the simulations where normal male subjects perform a long aerobic exercise following an overnight fast. These simulations replicate the experiments reported in [79] and [80]. Ahlborg and Felig [79] observed 20 normal male subjects as they performed a leg exercise at intensity 58 %*V O*_2_*max* for 3 to 3.5 hours after a 12 to 14 hour overnight fast. In a later study, Ahlborg et. al. [80] observed 12 normal male subjects as they performed a leg/arm exercise at intensity 30 %*V O*_2_*max* for 2 hours after a 12 to 14 hour overnight fast. The characteristics of these subjects are shown in Table 6. For each subject, the concentrations in the blood were recorded for a number of substrates and hormones including glucose. The relevant BGL data reported in [79] and [80] is shown in Table 7 and Fig 2. In the following subsections, we interpret each set of BGL data and demonstrate that with proper configuration *CarbMetSim* can be made to replicate each pattern.

**Table 6.**
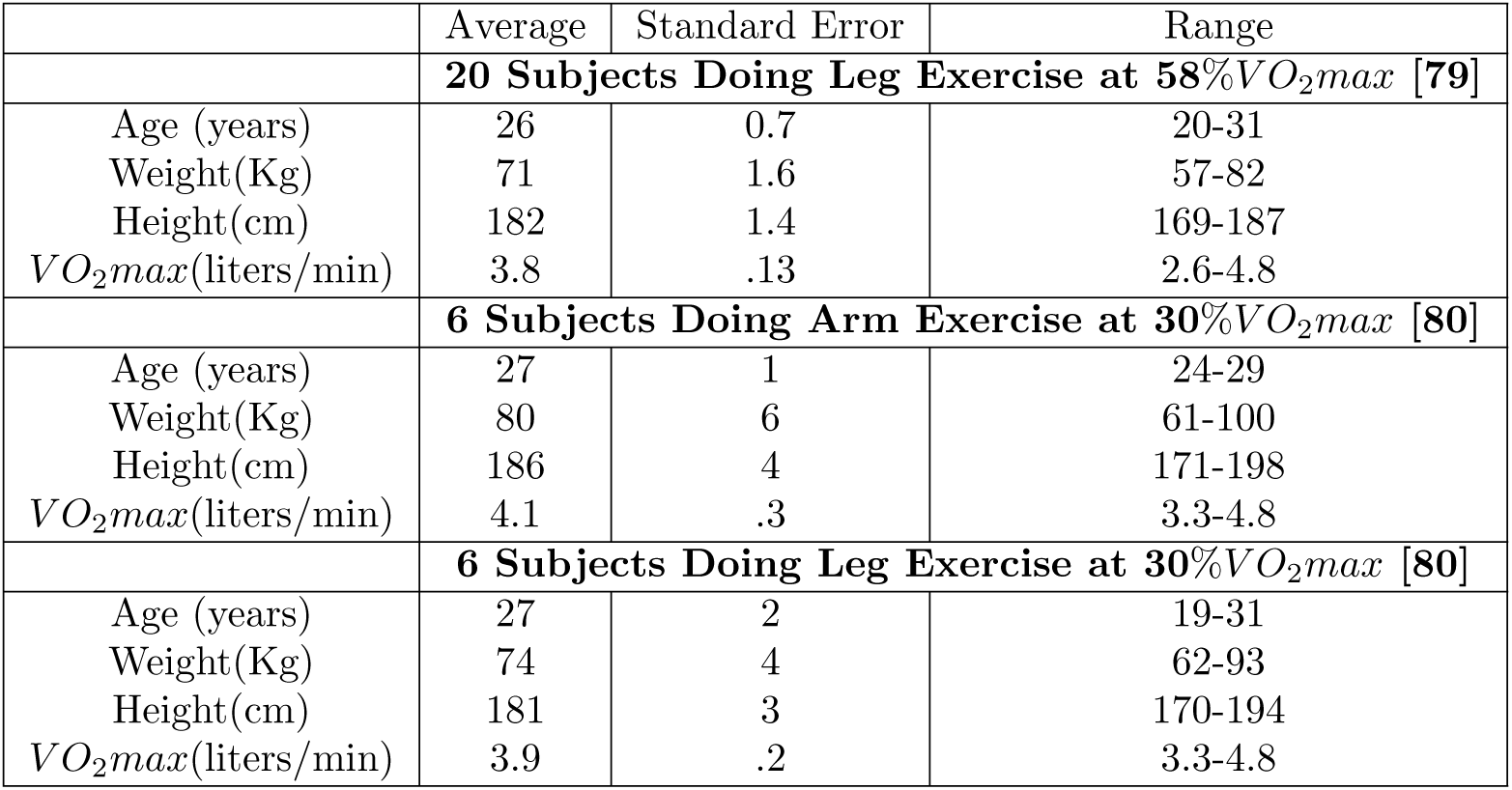
Characteristics of Subjects Reported in [79] and [80].

**Table 7.** BGL Measurements Reported in [79] and [80].

| | Blood Glucose Level (mmol/l): Average ± Standard Error | | |
| --- | --- | --- | --- |
| | Leg Exercise at 58%$VO_2max$ [79] | Arm Exercise at 30%$VO_2max$ [80] | Leg Exercise at 30%$VO_2max$ [80] |
| Rest | 4.39±0.08 | 4.00±0.11 | 4.33±0.09 |
| Exercise:40min | 4.09±0.10 | 4.01±0.31 | 4.28±0.10 |
| Exercise:90min | 3.86±0.28 | 4.06±0.20 | 4.07±0.16 |
| Exercise:120min | 3.55±0.11 | 3.98±0.23 | 3.81±0.15 |
| Exercise:180min | 2.78±0.13 | | |
| Exercise:210min | 2.56±0.13 | | |
| Recovery:10min | 3.12±0.13 | 3.96±0.31 | 4.06±0.25 |
| Recovery:20min | 3.19±0.13 | 3.76±0.29 | 4.11±0.25 |
| Recovery:40min | 3.18±0.10 | 3.83±0.25 | 4.13±0.21 |

**Fig 2.**
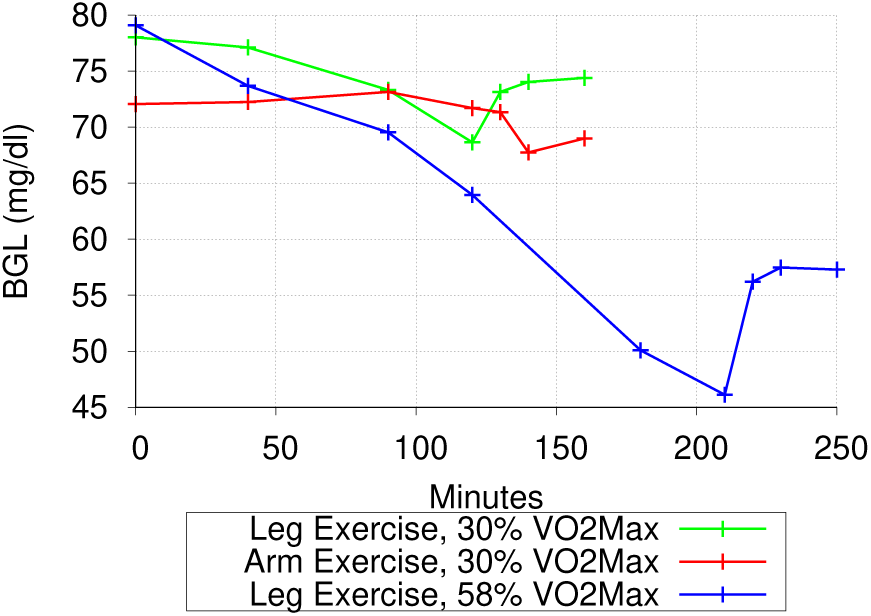
Average BGL Measurements (After Conversion to mg/dl) Reported in [79] and [80].

### 6.1 Exercise at Intensity 58 %*V O*_2_*max*

In the case of experiments involving physical exercise at intensity 58 %*V O*_2_*max*, Table 7 and Fig 2 show that there is a continuous drop in the BGL as the exercise progresses. The BGL approaches hypoglycemic levels towards the end of the exercise. Also, there is only a modest recovery from hypoglycemic BGL once the exercise concludes. These observations, coupled with the fact that the exercise began after a long fast, indicate that the liver glycogen was exhausted some time after the start of the exercise and that the local glycogen and the gluconeogenesis were the only sources of glucose for the exercising muscles. The BGL dropped continuously because the gluconeogenesis alone was not sufficient to compensate for the absorption of glucose from the blood by the exercising muscles. After the completion of the exercise, gluconeogenesis continues to be the only source of glucose and is insufficient to bring the BGL to pre-exercise level.

We generated 20 age and weight value pairs for normal male subjects to simulate using the average and standard error values specified in [79] (see Table 6) and simulated the described experiment on these subjects. Each simulation started at simulated time 12am and had the subject perform a 210 minute long exercise (at intensity 58 %*V O*_2_*max*) starting at 12pm. Each simulation ended at simulated time 5pm and used the same seed value for the random number generation. We adjusted the simulation parameters so as to cause the liver glycogen exhaustion early on in the exercise and thus match the BGL trends reported in [79]. The simulation parameters (that differed from the default values) for the simulated subjects are shown in Table 8. In each simulation, the initial glycogen store in the *Liver* was set to 60 grams so that very little glycogen was left in the *Liver* by the time the exercise event began at 12pm. The *gngImpact*_ parameter was set to values between 13.2 and 15.5 so as to appropriately limit the glucose production via gluconeogenesis during the exercise event.

**Table 8.** Configuration parameters for simulations for a single exercise event at intensity 58 %*V O*_2_*max*.

| Subject # | 1 | 2 | 3 | 4 | 5 | 6 | 7 | 8 | 9 | 10 | 11 | 12 | 13 | 14 | 15 | 16 | 17 | 18 | 19 | 20 |
| --- | --- | --- | --- | --- | --- | --- | --- | --- | --- | --- | --- | --- | --- | --- | --- | --- | --- | --- | --- | --- |
| age_ (years) | 23 | 26 | 26 | 30 | 22 | 25 | 26 | 24 | 24 | 22 | 20 | 23 | 31 | 26 | 26 | 30 | 21 | 29 | 25 | 26 |
| gender_ (0=Male) | 0 |  |  |  |  |  |  |  |  |  |  |  |  |  |  |  |  |  |  |  |
| fitnessLevel_ (%ile) | 50 |  |  |  |  |  |  |  |  |  |  |  |  |  |  |  |  |  |  |  |
| bodyWeight_ (kg) | 57 | 63 | 59 | 78 | 60 | 71 | 75 | 76 | 72 | 64 | 74 | 82 | 71 | 62 | 64 | 70 | 78 | 65 | 65 | 74 |
| minGlucoseLevel_ (mg/dl) | 40 |  |  |  |  |  |  |  |  |  |  |  |  |  |  |  |  |  |  |  |
| baseGlucoseLevel_ (mg/dl) | 79 |  |  |  |  |  |  |  |  |  |  |  |  |  |  |  |  |  |  |  |
| highGlucoseLevel_ (mg/dl) | 145 |  |  |  |  |  |  |  |  |  |  |  |  |  |  |  |  |  |  |  |
| baseInsulinLevel_ | 0.001 |  |  |  |  |  |  |  |  |  |  |  |  |  |  |  |  |  |  |  |
| peakInsulinLevel_ | 1.0 |  |  |  |  |  |  |  |  |  |  |  |  |  |  |  |  |  |  |  |
| gngImpact_ | 15.5 | 15.1 | 15.35 | 13.2 | 15.35 | 14.75 | 14.5 | 14.5 | 14.7 | 15.1 | 14.55 | 14.3 | 13.5 | 15.25 | 15.1 | 13.5 | 14.35 | 15 | 15 | 14.6 |
| Initial Liver Glycogen (g) | 60.0 |  |  |  |  |  |  |  |  |  |  |  |  |  |  |  |  |  |  |  |

Fig 3 shows the results of *CarbMetSim* simulations replicating the physical exercise event at intensity 58 %*V O*_2_*max* as reported in [79]. The BGL values for each simulated subject, along with the average BGL values reported in [79], are shown in Fig 3a. As this figure shows, the BGLs in the simulations have a close match with the measurements reported in [79]. For all subjects, the BGL hovered around its pre-exercise level for some time once the exercise began and then dropped continuously throughout the exercise duration with final values being in the hypoglycemic range. After the conclusion of the exercise activity, the BGL recovered but failed to reach its pre-exercise level.

**Fig 3.**
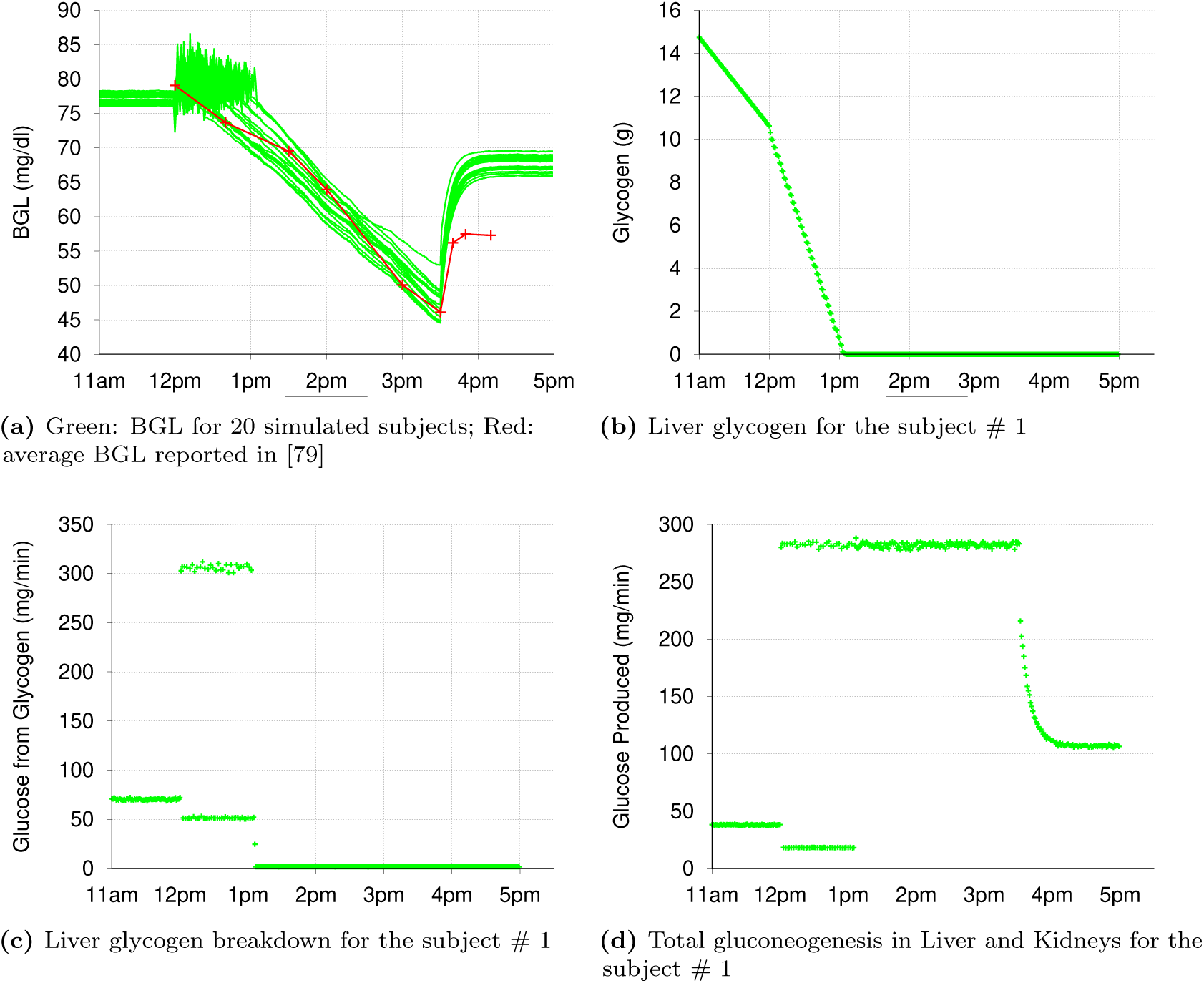
Results of simulations involving a physical exercise event at intensity 58 %*V O*_2_*max* as reported in [79].

Fig 3b shows the amount of glycogen left in the *Liver* for a particular simulated subject. As mentioned before, the initial amount of glycogen in the *Liver* was set so that all this glycogen would be exhausted in the early stages of the exercise activity. As is clear from Fig 3b, all the glycogen in the *Liver* was exhausted by 1pm after which the gluconeogenesis was the only source of glucose for this particular subject. Fig 3c and Fig 3d show the glycogen breakdown flux in the *Liver* and the combined gluconeogenesis flux in the *Liver* and *Kidneys* for this particular subject. Note that the glycogen breakdown flux and the gluconeogenesis flux fluctuated between high and low values once the exercise began (and before the *Liver* glycogen was exhausted). This behavior is in accordance with the manner in which the *insulinLevel* varies (Section 3.2) and the manner in which the glycogen breakdown in the *Liver* (Section 3.6) and gluconeogenesis in the *Liver* and *Kidneys* (Section 3.5) react to the *insulinLevel*. As described in Section 3.2, when the BGL falls below the *baseGlucoseLevel*_, the *insulinLevel* becomes zero if the body is engaged in an exercise at an intensity higher than the *intensityPeakGlucoseProd*_ (default value 20 %*V O*_2_*max*). Since the exercise intensity (58 %*V O*_2_*max*) was indeed higher than the *intensityPeakGlucoseProd*, the *insulinLevel* fell to zero whenever the BGL fell below the *baseGlucoseLevel*_. This caused both glycogen breakdown in the *Liver* and gluconeogenesis in the *Liver* and *Kidneys* to proceed at the highest levels. When the BGL exceeded the *baseGlucoseLevel*_, the *insulinLevel* exceeded the *baseInsulinLevel* and the glycogen breakdown & gluconeogenesis fluxes dropped down to the regular levels.

Once the liver glycogen was exhausted, the gluconeogenesis alone (even when occurring at the highest level) was not sufficient to push BGL above the *baseGlucoseLevel*_ and hence the *insulinLevel* stayed at zero level for rest of the exercise duration and the gluconeogenesis continued to occur at its highest level as the only source of glucose for the blood. Once the exercise was over, the *insulinLevel* increased to a positive value below *baseInsulinLevel*_ (as per the rules described in Section 3.2) and in response the gluconeogenesis flux assumed a value between the regular and the highest levels (as per the rules described in Section 3.5). Gluconeogenesis at this level allowed the BGL to climb up from the hypoglycemic range to a level below the *baseGlucoseLevel*_.

### 6.2 *Arm* Exercise at Intensity 30 %*V O*_2_*max*

In the case of experiments involving an *arm* exercise at intensity 30 %*V O*_2_*max*, it is clear from Table 7 and Fig 2 that the BGL largely maintains its pre-exercise level during the entire exercise duration (although there is a small drop in the BGL during the recovery phase). These observations indicate that the liver glycogen was not exhausted during the exercise and that the breakdown of liver glycogen and gluconeogenesis together were sufficient to meet the needs of the exercising muscles. Once the exercise was over, the liver glycogen breakdown and gluconeogenesis returned to their pre-exercise levels and accordingly the BGL also returned to its pre-exercise level.

We generated 6 age and weight value pairs for normal male subjects to simulate using the average and standard error values specified in [80] for the *arm* exercise experiments (see Table 6) and simulated the described experiment on these subjects. *CarbMetSim* does not currently distinguish between different muscles and hence the *arm* exercise was simulated as a regular exercise. Each simulation started at 12am and had the subject perform a 120 minute long exercise (at intensity 30 %*V O*_2_*max*) starting at 12pm. Each simulation ended at 5pm and used the same seed value for the random number generation. The simulation parameters (that differed from the default values) for the simulated subjects are shown in Table 9. In each simulation, the initial glycogen store in the *Liver* was set to 100 grams so that the liver glycogen does not get exhausted during the exercise. The *gngImpact*_ parameter was set to value 15.0 so that the glucose production via gluconeogenesis can ramp up to a high enough level when required during the exercise.

**Table 9.** Configuration parameters for simulations for a single “arm” exercise event at intensity 30 %*V O*_2_*max*.

| Subject # | 1 | 2 | 3 | 4 | 5 | 6 |
| --- | --- | --- | --- | --- | --- | --- |
| age_ (years) | 24 | 29 | 28 | 27 | 27 | 28 |
| gender_ (0=Male) | 0 |  |  |  |  |  |
| fitnessLevel_ (%ile) | 50 |  |  |  |  |  |
| bodyWeight (kg) | 61 | 100 | 97 | 62 | 88 | 87 |
| minGlucoseLevel_ (mg/dl) | 40 |  |  |  |  |  |
| baseGlucoseLevel_ (mg/dl) | 72 |  |  |  |  |  |
| highGlucoseLevel_ (mg/dl) | 145 |  |  |  |  |  |
| baseInsulinLevel_ | 0.001 |  |  |  |  |  |
| peakInsulinLevel_ | 1.0 |  |  |  |  |  |
| gngImpact_ | 15.0 |  |  |  |  |  |
| Initial Liver Glycogen (g) | 100.0 |  |  |  |  |  |

Fig 4 shows the results of *CarbMetSim* simulations replicating the *arm* exercise event at intensity 30 %*V O*_2_*max* as reported in [80]. The BGL values for each simulated subject, along with the average BGL values reported in [80], are shown in Fig 4a. As this figure shows, the BGLs in the simulations have a close match with the measurements reported in [80]. For all subjects, the BGL hovered around its pre-exercise level throughout the exercise duration and then went back to the pre-exercise level. Fig 4b shows the amount of glycogen left in the *Liver* for a particular simulated subject. As desired, the liver glycogen did not get exhausted during the exercise and in the recovery phase. Fig 4c and Fig 4d show the glycogen breakdown flux in the *Liver* and the combined gluconeogenesis flux in the *Liver* and *Kidneys* for this particular subject. As was the case with the 58 %*V O*_2_*max* simulations reported in the previous section, the glycogen breakdown flux and the gluconeogenesis flux fluctuated between high and low values once the exercise began. The explanation for this behavior was provided in the previous section. These oscillations explain the BGL oscillations throughout the exercise duration. Once the exercise was over, the *insulinLevel* increased to the *baseInsulinLevel*_ (as per the rules described in Section 3.2) and in response the liver glycogen breakdown and gluconeogenesis fluxes (and hence the BGL) assumed their pre-exercise levels.

**Fig 4.**
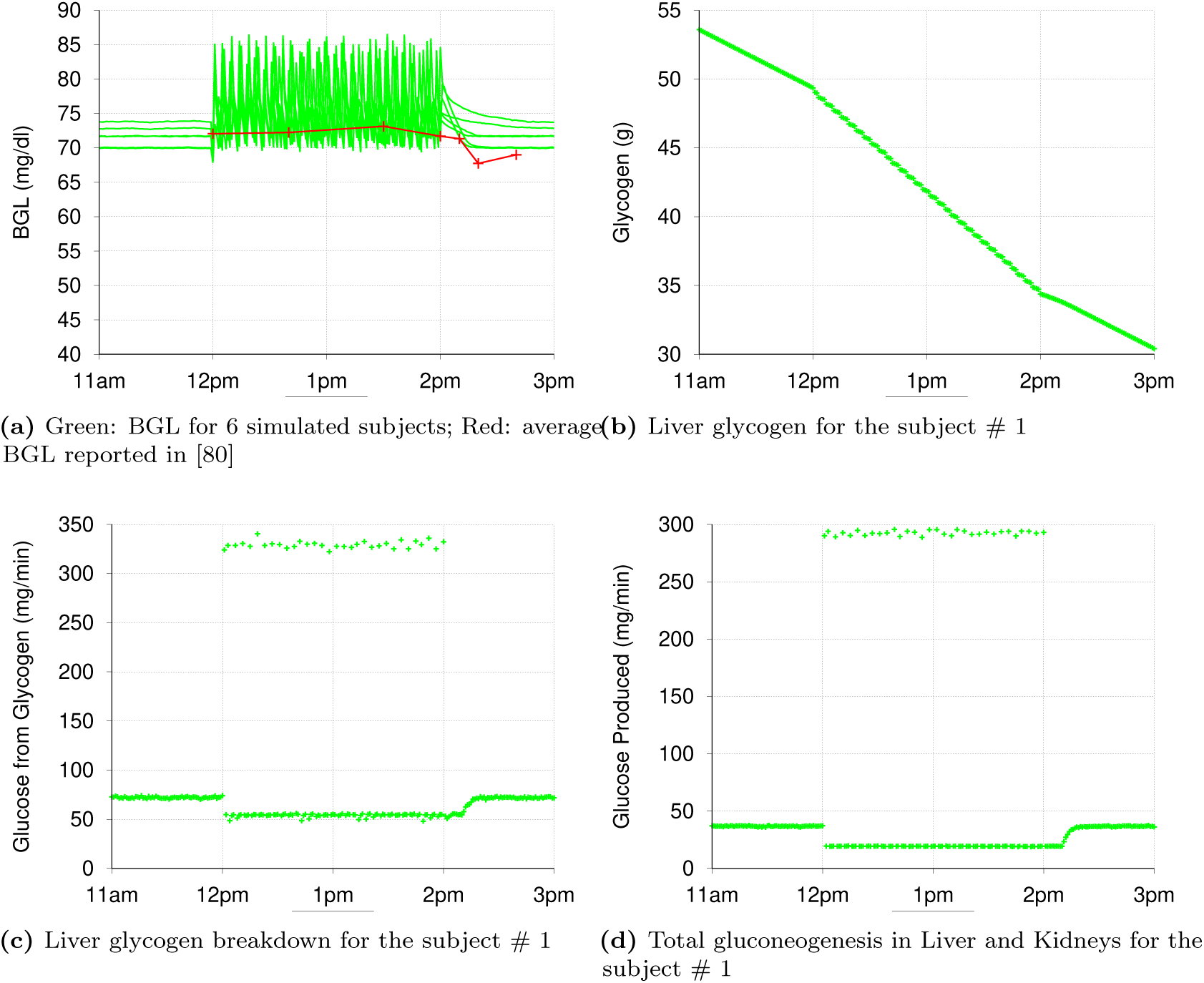
Results of simulations replicating a physical exercise event involving arms at intensity 30 %*V O*_2_*max* as reported in [80].

### 6.3 *Leg* Exercise at Intensity 30 %*V O*_2_*max*

Finally, we have the experiments involving a *leg* exercise at intensity 30 %*V O*_2_*max*. In these experiments, the BGL dropped modestly during the exercise and then seemed to climb back to the pre-exercise level (see Table 7 and Fig 2). The fact that the post-exercise BGL approached pre-exercise level indicates that the liver glycogen was not exhausted during the exercise or in the recovery phase. However, the fact that the BGL dropped modestly throughout the exercise indicates that glucose production during exercise (via liver glycogen breakdown and gluconeogenesis) was a little less than the amount absorbed from the blood by the exercising muscles. Clearly, in these experiments, the physical exercise was not able to stimulate liver glycogen breakdown and gluconeogenesis sufficiently so that their combined glucose production could match the demands of the exercising muscles.

We generated 6 age and weight value pairs for normal male subjects to simulate using the average and standard error values specified in [80] for the *leg* exercise experiments (see Table 6) and simulated the described experiment on these subjects. As mentioned before, *CarbMetSim* does not currently distinguish between different muscles and hence the *leg* exercise was simulated as a regular exercise. Each simulation started at 12am and had the subject perform a 120 minute long exercise (at intensity 30 %*V O*_2_*max*) starting at 12pm. Each simulation ended at 5pm and used the same seed value for the random number generation. In these simulations, we wanted to precisely control the glucose production via liver glycogen breakdown and gluconeogenesis during exercise. The total glucose production during exercise had to be just a little less than what the exercising muscles were absorbing from the blood. To achieve this end, we reduced the *liverGlycogenBreakdownImpact*_ (that controls the liver glycogen breakdown during exercise; see Sections 3.6 and 3.2) to value 1.0 (i.e. no extra glycogen breakdown in the *Liver* during the exercise) while increasing the *glycogenToGlucoseInLiver*_ parameter (that controls the regular glycogen breakdown in the *Liver*) value appropriately. The *gngImpact*_ parameter (that controls the gluconeogenesis flux during exercise; see Sections 3.5 and 3.2) was also set appropriately to limit glucose production via gluconeogenesis during exercise. All simulation parameters (that differed from the default values) for the simulated subjects are shown in Table 10. In each simulation, the initial glycogen store in the *Liver* was set to 100 grams so that the liver glycogen would not be exhausted during the exercise or the recovery phase.

**Table 10.** Configuration parameters for simulations for a single “leg” exercise event at intensity 30 %*V O*_2_*max*.

| Subject # | 1 | 2 | 3 | 4 | 5 | 6 |
| --- | --- | --- | --- | --- | --- | --- |
| age_ (years) | 20 | 31 | 22 | 29 | 30 | 25 |
| gender_ (0=Male) | 0 |  |  |  |  |  |
| fitnessLevel_ (%ile) | 50 |  |  |  |  |  |
| bodyWeight (kg) | 62 | 93 | 68 | 70 | 82 | 71 |
| minGlucoseLevel_ (mg/dl) | 40 |  |  |  |  |  |
| baseGlucoseLevel_ (mg/dl) | 78 |  |  |  |  |  |
| highGlucoseLevel_ (mg/dl) | 145 |  |  |  |  |  |
| baseInsulinLevel_ | 0.001 |  |  |  |  |  |
| peakInsulinLevel_ | 1.0 |  |  |  |  |  |
| gngImpact_ | 6.2 | 5.6 | 6.2 | 6.2 | 5.6 | 6.1 |
| Initial Liver Glycogen (g) | 100.0 |  |  |  |  |  |
| glycogenToGlucoseInLiver_ (mg/kg/min) | 1.4 | 0.9 | 1.3 | 1.25 | 1.05 | 1.25 |
| liverGlycogenBreakdownImpact_ | 1.0 |  |  |  |  |  |

Fig 5 shows the results of *CarbMetSim* simulations replicating the *leg* exercise event at intensity 30 %*V O*_2_*max* as reported in [80]. The BGL values for each simulated subject, along with the average BGL values reported in [80], are shown in Fig 5a. As this figure shows, the BGLs in the simulations have a reasonably good match with the measurements reported in [80]. As desired, for all subjects, the BGL dropped modestly during the exercise duration and then went back to the pre-exercise level. Fig 5b shows the amount of glycogen left in the *Liver* for a particular simulated subject and Fig 5c shows the glycogen breakdown flux in the *Liver* for this subject. As desired, the liver glycogen flux did not increase during the exercise. Fig 5d shows the combined gluconeogenesis flux in the *Liver* and *Kidneys* for this particular subject. Since the BGL was always below the *baseGlucoseLevel* throughout the exercise duration and the exercise intensity was greater than *intensityPeakGlucoseProd*_, the *insulinLevel* was zero throughout the exercise duration and hence the gluconeogenesis took place at its highest level throughout the exercise duration. However, the zero value of the *insulinLevel* was not able to stimulate liver glycogen breakdown because the *liverGlycogenBreakdownImpact*_ was set to value 1. The combined glucose production via gluconeogenesis and liver glycogen breakdown was just below the glucose absorbed from the blood by the exercising muscles and hence the BGL dropped modestly throughout the exercise duration as desired. Once the exercise was over, the *insulinLevel* increased to the *baseInsulinLevel*_ and in response the gluconeogenesis flux (and hence the BGL) assumed their pre-exercise level.

**Fig 5.**
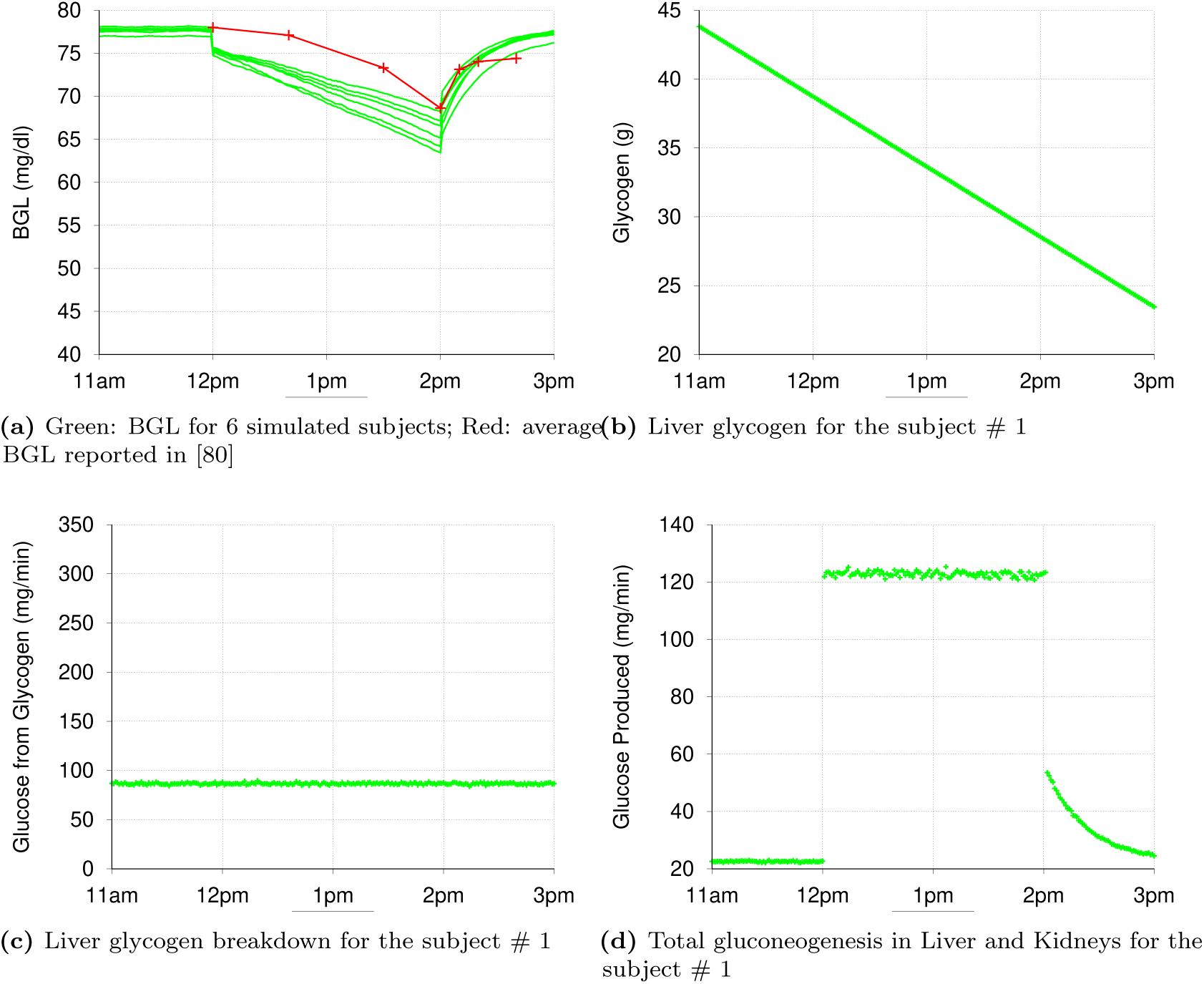
Results of simulations replicating a physical exercise event involving legs at intensity 30 %*V O*_2_*max* as reported in [80].

## 7 Conclusion

This paper described *CarbMetSim*, a discrete event simulator that models the carbohydrate metabolism in human beings and allows tracking of a normal or Type 2 Diabetic subject’s BGL in response to a timed sequence of diet and exercise activities. The paper also validated *CarbMetSim*’s behavior in response to single meal and exercise events and demonstrated its ability to emulate actual BGL patterns with appropriate configuration. Our future work on *CarbMetSim* will include more validation of its behavior against real BGL data, expanding its functionality to correct some of its current limitations identified towards the end of Section 1 and building web/smartphone apps that will allow diabetes patients to use the simulator. *CarbMetSim* can also serve as the underlying engine for a variety of diabetes self management and education tools. With its open source nature and ease of modification/extension, *CarbMetSim* also has a good potential to emerge as a popular simulation framework for diabetes research.

## References

1. International Diabetes Federation. IDF Diabetes Atlas - 8th Edition; 2017. http://www.diabetesatlas.org/

2. Kovatchev B, Breton M, Dalla Man C, Cobelli C. In Silico Preclinical Trials: A Proof of Concept in Closed-loop Control of Type 1 Diabetes. J Diabetes Sci Technol. 2009;3(1):44–55.

3. Dalla Man C, Micheletto F, Lv D, Breton M, Kovatchev B, Cobelli C. The UVA/PADOVA Type 1 Diabetes Simulator: New Features. J Diabetes Sci Technol. 2014;8(1):26–34.

4. Goyal M, Aydas B, Ghazaleh H. CarbMetSim: The Carbohydrate Metabolism Simulator; 2018. https://github.com/mukulgoyalmke/CarbMetSim

5. Cobelli C, Dalla Man C, Sparacino G, Magni L, De Nicolao G, Kovatchev BP. Diabetes: Models, Signals, and Control. IEEE Rev Biomed Eng. 2009;2:54–96.

6. Tresp V, Briegel T, Moody J. Neural-network Models for the Blood Glucose Metabolism of a Diabetic. IEEE Trans Neural Netw. 1999;10(5):1204–1213.

7. Zitar RA, Al-Jabali A. Towards Neural Network Model for Insulin/Glucose in Diabetics-II. Informatica. 2005;29:227–232.

8. Quchani S, Tahami E. Comparison of MLP and Elman Neural Network for Blood Glucose Level Prediction in Type 1 Diabetics. In: 3rd Kuala Lumpur International Conference on Biomedical Engineering 2006. Springer; 2006. p. 54–58.

9. Baghdadi G, Nasrabadi AM. Controlling Blood Glucose Levels in Diabetics by Neural Network Predictor. In: 29th Annual International Conference of the IEEE Engineering in Medicine and Biology Society. IEEE; 2007. p. 3216–3219.

10. Zainuddin Z, Pauline O, Ardil C. A Neural Network Approach in Predicting the Blood Glucose Level for Diabetic Patients. Int J Comput Intell. 2009;5(1):72–79.

11. Mougiakakou SG, Prountzou A, Iliopoulou D, Nikita KS, Vazeou A, Bartsocas CS. Neural Network Based Glucose-insulin Metabolism Models for Children with Type 1 Diabetes. In: 28th Annual International Conference of the IEEE Engineering in Medicine and Biology Society. IEEE; 2006. p. 3545–3548.

12. Valletta JJ, Chipperfield AJ, Byrne CD. Gaussian Process Modelling of Blood Glucose Response to Free-living Physical Activity Data in People with Type 1 Diabetes. In: Annual International Conference of the IEEE Engineering in Medicine and Biology Society. IEEE; 2009. p. 4913–4916.

13. Oviedo S, Vehí J, Calm R, Armengol J. A Review of Personalized Blood Glucose Prediction Strategies for T1DM Patients. International journal for numerical methods in biomedical engineering. 2017;33(6).

14. Boliei V. Coefficients of Normal Blood Glucose Regulation. J Clin Invest. 1960;39(2):783–788.

15. Ackerman E, Gatewood L, Rosevear J, Molnar G. Model Studies of Blood-glucose Regulation. Bulletin of Math Biophy. 1965;27:21–37.

16. Gatewood L, Ackerman E, Rosevear J, Molnar G. Simulation Studies of Blood-glucose Regulation: Effect of Intestinal Glucose Absorption. Comput and Biomed Research. 1968;2:15–27.

17. Charrette W. Control System Theory Applied to Metabolic Homeostatic Systems and the Derivation and Identification of Mathematical Models. California Institute of Technology; 1968.

18. Foster R, Soeldner J, Tan M, Guyton J. Short Term Glucose Homeostasis in Man: A Systems Dynamic Model. J Dyn Syst Meas Control. 1973;95:308–314.

19. Cerasi E, Fick G, Rudemo M. A Mathematical Model for the Glucose Induced Insulin Release in Man. Europ J Clin Invest. 1974;4:267–278.

20. Insel P, Liljenquist J, Tobin J, Sherwin R, Watkins P, Andres R, et al. Insulin Control of Glucose Metabolism in Man. J Clin Invest. 1975;55:1057–1066.

21. Cramp D, Carson E. The Dynamics of Short-term Blood Glucose Regulation. In: Carbohydrate Metabolism. John Wiley and Sons; 1981. p. 349–368.

22. Cobelli C, Federspil G, Pacini G, Salvan A, Scandellari C. An Integrated Mathematical Model of the Dynamics of Blood Glucose and its Hormonal Control. Mathematical Biosciences. 1982;58:27–60.

23. Sorensen J. A Physiologic Model of Glucose Metabolism in Man and its Use to Design and Assess Improved Insulin Therapies for Diabetes. Massachusetts Institute of Technology; 1978.

24. Bergman RN, Ider YZ, Bowden CR, Cobelli C. Quantitative Estimation of Insulin Sensitivity. American Journal of Physiology-Endocrinology And Metabolism. 1979;236(6):E667–E677.

25. Bergman RN, Phillips LS, Cobelli C. Physiologic Evaluation of Factors Controlling Glucose Tolerance in Man: Measurement of Insulin Sensitivity and Beta-cell Glucose Sensitivity from the Response to Intravenous Glucose. Journal of clinical investigation. 1981;68(6):1456–1467.

26. Furler SM, Kraegen EW, Smallwood RH, Chisholm DJ. Blood Glucose Control by Intermittent Loop Closure in the Basal Mode: Computer Simulation Studies with a Diabetic Model. Diabetes Care. 1985;8(6):553–561.

27. Ollerton R. Application of Optimal Control Theory to Diabetes Mellitus. International Journal of Control. 1989;50(6):2503–2522.

28. Fisher ME. A Semiclosed-loop Algorithm for the Control of Blood Glucose Levels in Diabetics. IEEE Trans Biomed Eng. 1991;38(1):57–61.

29. Roy A, Parker RS. Dynamic Modeling of Free Fatty Acid, Glucose, and Insulin: An Extended “Minimal Model”. Diabetes Technology & Therapeutics. 2006;8(6):617–626.

30. Roy A, Parker RS. Dynamic Modeling of Exercise Effects On Plasma Glucose and Insulin Levels. IFAC Proceedings Volumes. 2006;39(2):509–514.

31. Derouich M, Boutayeb A. The Effect of Physical Exercise on the Dynamics of Glucose and Insulin. Journal of Biomechanics. 2002;35(7):911–917.

32. Tiran J, Avruch I, Albisser A. A Circulation and Organs Model for Insulin Dynamics. Am J Physiol. 1979;237:E331–E339.

33. Guyton JR, Foster RO, Soeldner JS, Tan MH, Kahn CB, Koncz L, et al. A Model of Glucose-insulin Homeostasis in Man that Incorporates the Heterogeneous Fast Pool Theory of Pancreatic Insulin Release. Diabetes. 1978;27(10):1027–1042.

34. Parker RS, Doyle FJ, Peppas NA. A Model-based Algorithm for Blood Glucose Control in Type I Diabetic Patients. IEEE Trans Biomed Eng. 1999;46(2):148–157.

35. Parker RS, Doyle FJ, Ward JH, Peppas NA. Robust H*∞* Glucose Control in Diabetes using a Physiological Model. AIChE Journal. 2000;46(12):2537–2549.

36. Lehmann E, Deutsch T. A Physiological Model of Glucose-insulin Interaction in Type I Diabetes Mellitus. J Biomed Eng. 1992;14:235–242.

37. Hovorka R, Canonico V, Chassin LJ, Haueter U, Massi-Benedetti M, Federici MO, et al. Nonlinear Model Predictive Control of Glucose Concentration in Subjects with Type 1 Diabetes. Physiological Measurement. 2004;25:905–920.

38. Dalla Man C, Rizza RA, Cobelli C. Meal Simulation Model of the Glucose-insulin System. IEEE Trans Biomed Eng. 2007;54(10):1740–1749.

39. Englyst HN, Kingman S, Cummings J. Classification and Measurement of Nutritionally Important Starch Fractions. European Journal of Clinical Nutrition. 1992;46:S33–50.

40. Frayn K. Metabolic Regulation: A Human Perspective, 3rd Edition. Wiley-Blackwell; 2010.

41. Gleeson M. Interrelationship between Physical Activity and Branched-Chain Amino Acids. J Nutrition. 2005;135(6):1591S–1595S.

42. Ploug T, Galbo H, Richter E. Increased Muscle Glucose Uptake During Contractions: No Need for Insulin. American Journal of Physiology. 1984;247:E726–E731.

43. Goodyear L, King P, Hirshman M, Thompson C, Horton E. Contractile Activity Increases Plasma Membrane Glucose Transporters in Absence of Insulin. American Journal of Physiology. 1990;258:E667–E672.

44. Wasserman D, Zinman B. Exercise in Individuals with IDDM. Diabetes Care. 1994;17(8):924–937.

45. Berger M, Berchtold P, Cuppers H, Drost H, Kley H, Muller W, et al. Metabolic and Hormonal Effects of Muscular Exercise in Juvenile Type Diabetics. Diabetologia. 1977;13:355–365.

46. Kemmer F, Berchtold P, Berger M, Starke A, Cuppers HJ, Gries F, et al. Exercise-induced Fall of Blood Glucose in Insulin-treated Diabetics Unrelated to Alteration of Insulin Mobilization. Diabetes. 1979;28:1131–1137.

47. Horton E. Role and Management of Exercise in Diabetes Mellitus. Diabetes Care. 1988;11(2):201–211.

48. Meyer C, Stumvoll M, Dostou J, Welle S, Haymond M, Gerich J. Renal Substrate Exchange and Gluconeogenesis in Normal Postabsorptive Humans. Am J Physiol Endocrinol Metab. 2002;282:E428–E434.

49. Magnusson I, Rothman D, Katz L, Shulman R, Shulman G. Increased Rate of Gluconeogenesis in Type II Diabetes Mellitus. A 13C Nuclear Magnetic Resonance Study. Journal of Clinical Investigation. 1992;90(4):1323.

50. Puhakainen I, Koivisto V, Yki-Jarvinen H. Lipolysis and Gluconeogenesis from Glycerol are Increased in Patients with Noninsulin-dependent Diabetes Mellitus. J Clin Endocrinol Metab. 1992;75(3):789–794.

51. Woerle H, Szoke E, Meyer C, Dostou J, Wittlin S, Gosmanov N, et al. Mechanisms for Abnormal Postprandial Glucose Metabolism in Type 2 Diabetes. Am J Physiol Endocrinol Metab. 2006;290:E67–E77.

52. Gerich J. Role of the Kidney in Normal Glucose Homeostasis and in the Hyperglycaemia of Diabetes Mellitus: Therapeutic Implications. Diabetic Medicine. 2010;27(2):136–142.

53. Woerle HJ, Meyer C, Dostou JM, Gosmanov NR, Islam N, Popa E, et al. Pathways for Glucose Disposal after Meal Ingestion in Humans. American Journal of Physiology-Endocrinology and Metabolism. 2003;284(4):E716–E725.

54. Kaminsky L, Arena R, Myers J. Reference Standards for Cardiorespiratory Fitness Measured with Cardiopulmonary Exercise Testing: Data From the Fitness Registry and the Importance of Exercise National Database. Mayo Clinic Proceedings. 2015;90(11):1515–1523.

55. Low A. Nutritional Regulation of Gastric Secretion, Digestion and Emptying. Nutrition Research Reviews. 1990;3:229–252.

56. Minami H, McCallum R. The Physiology and Pathophysiology of Gastric Emptying in Humans. Gastroenterology. 1984;86:1592–1610.

57. Hunt J, Smith J, Jiang C. Effect of Meal Volume and Energy Density on the Gastric Emptying of Carbohydrates. Gastroenterology. 1985;89:1326–1330.

58. McHugh P, Moran T. Calories and Gastric Emptying: A Regulatory Capacity with Implications for Feeding. Am J Physiol. 1979;236:R254–R260.

59. Hunt J, Spurrell W. The Pattern of Emptying of the Human Stomach. J Physiol. 1951;113:157–168.

60. Elashoff J, Reedy T, Meyer J. Analysis of Gastric Emptying Data. Gastroenterology. 1982;83:1306–1312.

61. Dalla Man C, Camilleri M, Cobelli C. A System Model of Oral Glucose Absorption: Validation on Gold Standard Data. IEEE Trans Biomed Eng. 2006;53(12):2472–2478.

62. Hunt J, Stubbs D. The Volume and Energy Content of Meals as Determinants of Gastric Emptying. J Physiol. 1975;245:209–225.

63. Romijn J, Coyle E, Sidossis L, Gastaldelli A, Horowitz J, Endert E, et al. Regulation of Endogenous Fat and Carbohydrate Metabolism in Relation to Exercise Intensity and Duration. American Journal of Physiology. 1993;265:E380–E391.

64. Moe O, Wright S, Palacin M. Renal Handling of Organic Solutes. In: Brenner & Rector’s The Kidney, 8th edition. Saunders Elsevier; 2008. p. 214–247.

65. Ahlborg G, Felig P, Hagenfeldt L, Hendler R, Wahren J. Substrate Turnover During Prolonged Exercise in Man. Journal of Clinical Investigation. 1974;53:1080–1090.

66. Kelley D, Mitrakou A, Marsh H, Schwenk F, Benn J, Sonneberg G, et al. Skeletal Muscle Glycolysis, Oxidation, and Storage of an Oral Glucose Load. J Clin Invest. 1988;81:1563–1571.

67. Felig P, Wahren J. Fuel Homeostasis in Exercise. New England Journal of Medicine. 1975;293:1078–1084.

68. Abel E. Glucose Transport in the Heart. Front Biosci. 2004;9:201–215.

69. Nelson J, Poussier P, Marliss E, Albisser A, Zinman B. Metabolic Response of Normal Man and Insulin-infused Diabetics to Postprandial Exercise. American Journal of Physiology. 1982;242:E309–E316.

70. Caron D, Poussier P, Marliss E, Zinman B. The Effect of Postprandial Exercise on Meal-related Glucose Intolerance in Insulin-dependent Diabetic Individuals. Diabetes Care. 1982;5:364–369.

71. Larsen J, Dela F, Kjaer M, Galbo H. The Effect of Moderate Exercise on Postprandial Glucose Homeostasis in NIDDM Patients. Diabetologia. 1997;40:447–453.

72. Larsen J, Dela F, Madsbad S, Galbo H. The Effect of Intense Exercise on Postprandial Glucose Homeostasis in Type 2 Diabetic Patients. Diabetologia. 1999;42:1282–1292.

73. Soman V, Koivisto V, Deibert D, Felig P, DeFronzo R. Increased Insulin Sensivity and Insulin Binding to Monocytes After Physical Training. New England Journal of Medicine. 1979;301:1200–1204.

74. Wahren J, Felig P, Ahlborg G, Jorfeldt L. Glucose Metabolism During Leg Exercise in Man. Journal of Clinical Investigation. 1971;50:2715–2725.

75. Galbo H. The Hormonal Response to Exercise. Diabetes Metabolism Review. 1986;1:385–408.

76. Wolfe R, Nadel E, Shaw J, Stephenson L, Wolfe M. Role of Changes in Insulin and Glucagon in Glucose Homeostasis in Exercise. Journal of Clinical Investigation. 1986;77:900–907.

77. Kemmer F, Vranic M. The Role of Glucagon and its Relationship to Other Glucoregulatory Hormones in Exercise. In: Glucagon, Physiology, Pathophysiology and Morphology of the Pancreatic A-Cells. Elsevier; 1981. p. 297–331.

78. Zinman B, Murray F, Vranic M, Albisser A, Leibel B, McClean P, et al. Glucoregulation During Moderate Exercise in Insulin-treated Diabetics. Journal of Clinical Endocrinology and Metabolism. 1977;45:641–652.

79. Ahlborg G, Felig P. Lactate and Glucose Exchange across the Forearm, Legs, and Splanchnic Bed during and after Prolonged Leg Exercise. Journal of Clinical Investigation. 1982;69:45–54.

80. Ahlborg G, Wahren J, Felig P. Splanchnic and Peripheral Glucose and Lactate Metabolism during and after Prolonged Arm Exercise. Journal of Clinical Investigation. 1986;77:690–699.

